# Regulatory Mechanisms of Maternal Imprinting at the Dlk1-Dio3 Domain

**DOI:** 10.64898/2026.01.10.693661

**Authors:** Ariella Weinberg-Shukron, Fran L. Dearden, Ana Moreno-Barriga, Lok Ting Nick New, Marco Müller, Rebecca C. Rancourt, Carol A. Edwards, Anne C. Ferguson-Smith

## Abstract

Genomic imprinting is an epigenetic process causing parent-of-origin specific gene expression. The Dlk1-Dio3 domain is one of the largest imprinted clusters. While DNA methylation at an intergenic CpG-island (IG-CGI) within the imprinting control region (ICR) controls expression from the paternal chromosome, mechanisms regulating the unmethylated maternal chromosome remain unknown.

Within the transcriptional regulatory element (IG-TRE) of the ICR, deletions identified a minimal region *in vitro* exhibiting both silencing and enhancing activity, with SOX2 and ZFP281 contributing to enhancer function on the maternal chromosome.

*In vivo*, however, this deletion did not affect maternal expression in embryos; instead it activated *Dlk1* on both parental chromosomes. Combining deletion of this IG-TRE with the lethal IG-CGI deletion rescued lethality by balancing *Dlk1* expression, despite persistent maternal gene upregulation.

These results demonstrate that loss of expression at this domain is more detrimental than gain highlighting the importance of *in vivo* analysis. Identification of active regulatory factors on the unmethylated maternal chromosome challenges the prevailing view that imprinting is primarily a methylation-driven phenomenon, further revealing the sophisticated hierarchical mechanisms governing imprinting control.

## Introduction

Genome function is regulated temporally and tissue-specifically through the orchestrated interplay of regulatory factors, genomic features, and epigenetic states. Epigenetic modifications are dynamic during development and across the cell cycle. A hierarchy of successive epigenetic states, including DNA methylation, ensures the creation of healthy individuals. In mammals, extensive epigenetic reprogramming events occur during germ cell development, fertilization, and early embryogenesis^1^. While DNA methylation is essential for normal mammalian development, there appear to be multiple ways in which it can regulate and maintain cell fate and function.

One major process in which causal relationships between DNA methylation and gene expression is more comprehensively understood is that of mammalian genomic imprinting^2,3^. Genomic imprinting is an epigenetically regulated process causing genes to be expressed according to parent-of-origin. Imprinting is highly conserved in eutherian mammals, with mouse studies providing insights into the repertoire of developmental and physiological pathways regulated by imprinted genes^4,5^.

Differentially DNA methylated regions (DMRs) at imprinting control regions (ICRs) are established in the parental germlines^6^. They are protected from the global wave of demethylation in the early embryo by the KRAB zinc finger proteins ZFP57 and ZFP445 in order to maintain the epigenetic memory of parental origin^7^. Failure to correctly establish and maintain imprints is associated with developmental syndromes including growth abnormalities, neurological and metabolic disorders and numerous forms of cancer^8,9^. The extreme dosage sensitivity of imprinted genes highlights the complex and dynamic nature of their gene expression and importance in human biology.

Genomic imprinting in the laboratory mouse is a tractable model for studying epigenetic regulation, as the two genetically identical parental genomes within the same nucleus express a different repertoire of genes according to their parental origin. However, the dynamic hierarchy of events initiated by the ICR, which leads to the long-range, domain-wide, temporal and tissue-specific behaviour of imprinted genes, is not fully understood. By employing a systemic set of mutants at a single imprinted domain we can dissect the relationship between allelic expression, dosage, epigenetic control, and phenotypic outcomes.

The developmentally regulated *Dlk1-Dio3* imprinted domain, conserved between mouse and human, spans 1.2Mb and is one of the largest imprinted clusters. Failure to correctly imprint genes in this cluster in mice causes prenatal lethality, and in humans leads to Temple or Kagami-Ogata Syndromes, both exhibiting neurological, developmental and behavioural impairments^10,11^.

Four protein-coding genes: *Dlk1*, *Rtl1*, *Dio3* and one isoform of *Begain* are preferentially expressed from the paternally inherited chromosome and the maternally inherited chromosome expresses multiple imprinted noncoding transcripts, including *Gtl2* (also known as *Meg3*), *Rian* and *Mirg* (Fig. 1A)^12^. *Gtl2* also forms a long polycistronic transcript along with its associated transcripts *Rian* and *Mirg* acting as a host for multiple snoRNAs and miRNAs including the miR-379/miR-410 cluster, all of which are driven by the *Gtl2* promoter^13–15^. Appropriate expression of genes in this region is essential for the lifelong health of mammals and understanding their epigenetic regulation has significant biomedical relevance for a diverse range of processes ^16,17^.

**Figure 1-.**
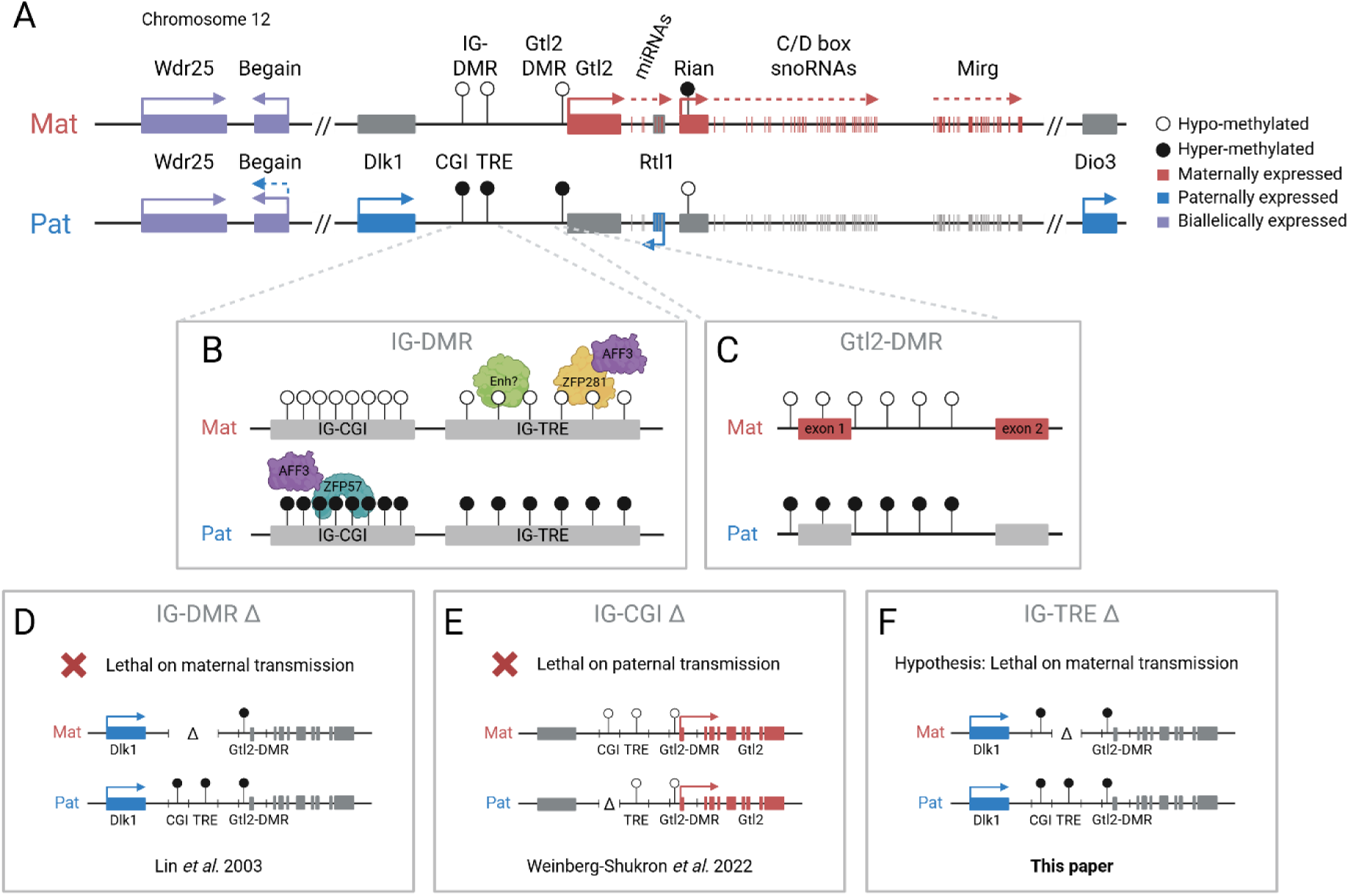
The *Dlk1-Dio3* imprinted domain, key regulatory features and deletion models: **A**. The paternally inherited chromosome expresses *Dlk1, Rtl1, Dio3* and one isoform of *Begain*. The maternally inherited chromosome expresses *Gtl2/Meg3, antiRtl1* and arrays of snoRNAs and miRNAs. The IG-DMR is methylated in sperm and unmethylated in oocytes and is the imprinting control region for the domain. DMRs are indicated by circles: black (methylated), white (unmethylated). **B**. The IG-DMR contains a CpG island (CGI) that binds ZFP57 (blue) on the methylated paternal copy and a transcriptional regulatory element (TRE) that has an enhancer-like function on the unmethylated maternal copy (green). The IG-DMR also binds AFF3 (purple) though ZFP57 at the IG-CGI on the methylated paternal chromosome and co-bound with ZFP281 (yellow) at the IG-TRE on the unmethylated maternal chromosome. **C**. The secondary somatic Gtl2-DMR is methylated *in cis* according to the methylation pattern of the IG-DMR. **D**. Maternal deletion of the entire IG-DMR results in a maternal-to-paternal epigenotype switch (Lin *et al*. 2003). **E**. Paternal deletion of the IG-CGI results in a paternal-to-maternal epigenotype switch (Weinberg-Shukron *et al*. 2022). **F**. Maternal deletion of the IG-TRE is hypothesized to result in a maternal-to-paternal epigenotype switch, similar to deletion of the entire IG-DMR.

Imprinting at the *Dlk1-Dio3* domain is regulated by the germline derived intergenic DMR (IG-DMR)^18^ which spans approximately 5kb between *Dlk1* and *Gtl2* and is required for regulating parent-specific expression across the locus^19^. This DMR is normally methylated in sperm and unmethylated in oocytes and maintains this parent-specific pattern throughout development (Fig. 1A,B). After implantation, a secondary somatic DMR is established at the promoter of the *Gtl2* gene (the Gtl2-DMR), which is also hypermethylated on the paternal chromosome^20^ (Fig. 1C).

The IG-DMR was recently established as a bipartite element comprised of two distinct functional elements^21^ (Fig. 1B). The first, termed the IG-CGI, is a small CpG island comprising seven tandemly repeated sequences of which five contain a ZFP57 binding motif^22^. The sequence-specific zinc finger protein, ZFP57, only binds to the methylated paternal chromosome, where it interacts with TRIM28 which recruits the repressive epigenetic machinery including DNA methyltransferases (DNMTs) and heterochromatin associated proteins, thereby maintaining the methylation memory of the germline imprint in early development at the time when most epigenetic modifications are being erased^23,24^. The IG-DMR has also been shown to bind AFF3, a component of the super elongation complex-like 3 (SEC-L3), on the methylated allele in mouse embryonic stem cells (mESCs) where it is thought to interact with the ZFP57/TRIM28 complex^25^ (Fig. 1B). However, the function of this interaction is unclear since it also binds a distal enhancer on the maternal chromosome, and depletion of AFF3 in mESCs leads to decreased expression of the maternally expressed genes.

Maternal deletion of the IG-DMR leads to paternalization of the maternal chromosome and mice die between e16.5 and birth^18,26^. However, on paternal transmission this deletion has no effect (Fig. 1D). Surprisingly, in contrast to the full IG-DMR deletion, paternal deletion of only the IG-CGI results in the reciprocal paternal-to-maternal epigenotype switch with the Gtl2-DMR becoming hypomethylated on the paternal chromosome^27,28^ (Fig. 1E) and maternal deletion of the IG-CGI has no effect. These data indicate that the IG-CGI is required to maintain a repressive chromatin landscape on the paternal chromosome, while the same element is dispensable on the maternal copy^28–30^.

The experiments therefore indicate that the maternal epigenotype must be regulated by elements within the IG-DMR, independent of the CpG island. Together, the paradoxical effects imposed by distinct deletions within the IG-DMR represent an attractive experimental framework for dissecting the impact of changes in gene dosage on embryonic phenotypes.

The remaining part of the IG-DMR, located downstream of the IG-CGI, has the potential to be an enhancer. It binds pluripotency transcription factors in mESCs, exhibiting active enhancer marks (H3K27ac) and nascent transcription^25,31^. It was therefore termed the transcriptional regulatory element (IG-TRE) and suggested to drive expression of the maternally inherited genes within the domain (Fig. 1B)^21^.

The transcription factor AFF3 also binds to the 3’ side of the IG-DMR downstream of the IG-TRE. There, AFF3 is co-bound with the zinc finger protein 281 (ZFP281/ZNF281), a zinc finger protein that has previously been reported to act as both a transcriptional activator and a repressor^32^ (Fig. 1B). In the *Dlk1-Dio3* locus depletion of ZFP281 from mESCs leads to decreased AFF3 binding at this downstream region. And since depletion of AFF3 leads to decreased expression of *Gtl2*, *Mirg* and *Rian*, this suggests that the downstream bound region is relevant for AFF3 function and that it also acts as an enhancer for maternally expressed genes^33^. Whether this region is acting in concert with the IG-TRE to control expression of *Gtl2* and its associated transcripts remains to be established.

Recently, Hara *et al*. demonstrated that deletion of a 2.7kb segment containing the IG-TRE results in postnatal lethality when maternally inherited^34^. However, smaller deletions of the IG-TRE subregions, including the AFF3/ZFP281 binding site alone, did not produce any obvious phenotype, suggesting the involvement of multiple elements spanning the IG-TRE in regulation of the maternal epigenotype.

There is evidence suggesting a role for pluripotency-associated transcription factors such as OCT4, SOX2, and KLF4 in maintaining the unmethylated state of the maternal allele at imprints, potentially by forming a regulatory complex that promotes enhancer activity in pluripotent cells. For example, at the Igf2-H19 imprinted locus, SOX2 and OCT4 bind the H19-ICR, and preserve the hypomethylated status of the maternal allele^35,36^. Whether this is a feature of these factors at other imprinted loci such as the IG-DMR is useful to ascertain. Therefore, while the multiple transcription factor binding motifs suggest that the IG-TRE is most likely a positive regulator for *Gtl2*, it is possible that the IG-TRE also contains a silencer element that is capable of directly repressing the paternally expressed genes on the maternally inherited chromosome. Experiments dissecting this region further are necessary to tease apart these options (Fig. 1F).

Here, we defined the minimal region of activity at the IG-TRE *in vitro*, generated an *in vivo* knockout mouse model to dissect the function of the IG-TRE during mouse development and piece these together with the previous deletion models to understand how the parental-origin specific epigenetic landscapes, genetic elements and transcription are exquisitely coordinated to regulate genome function at this domain.

## Results

### Functional dissection of the IG-DMR in vitro

The IG-DMR knockout in mice demonstrated that it regulates the activity and repression of multiple imprinted genes *in cis* and influences differential methylation at the Gtl2-DMR in the Dlk1-Dio3 domain^18^ but, though studies have shown it contains two features (IG-CGI and IG-TRE) acting individually on the paternally and maternally inherited imprinted gene cluster respectively, a systematic dissection of the regulatory features within the IG-DMR has not been conducted making it unclear how it works. In order to address this, we used an *in vitro* luciferase reporter gene assay in a manner described for other imprinted clusters^37,38^.

A 6.125kb region of the IG-DMR was divided into four overlapping fragments (Fig. 2A). The first 2kb of the IG-DMR knockout region is referred to as the 5’ fragment, which encompasses the IG-CGI and contains tandem repeat arrays identified by Paulsen *et al*^39^. The next fragment, referred to as Mid, overlaps the 5’ fragment and is located in the core of the differentially methylated region. It is 1.2kb and also contains the CpG island and the tandem repeat array. At the end of the original knockout region^18^ is the 3’1 fragment which is 1.3kb and located within the differentially methylated region. Finally, the 3’2 fragment spanning 2.2kb includes the 3’ sequence downstream of the IG-DMR knockout region. All fragments were cloned in both orientations, with the sense (S) referring to the direction towards *Gtl2* and antisense (AS) referring to the direction towards *Dlk1*.

**Figure 2-.**
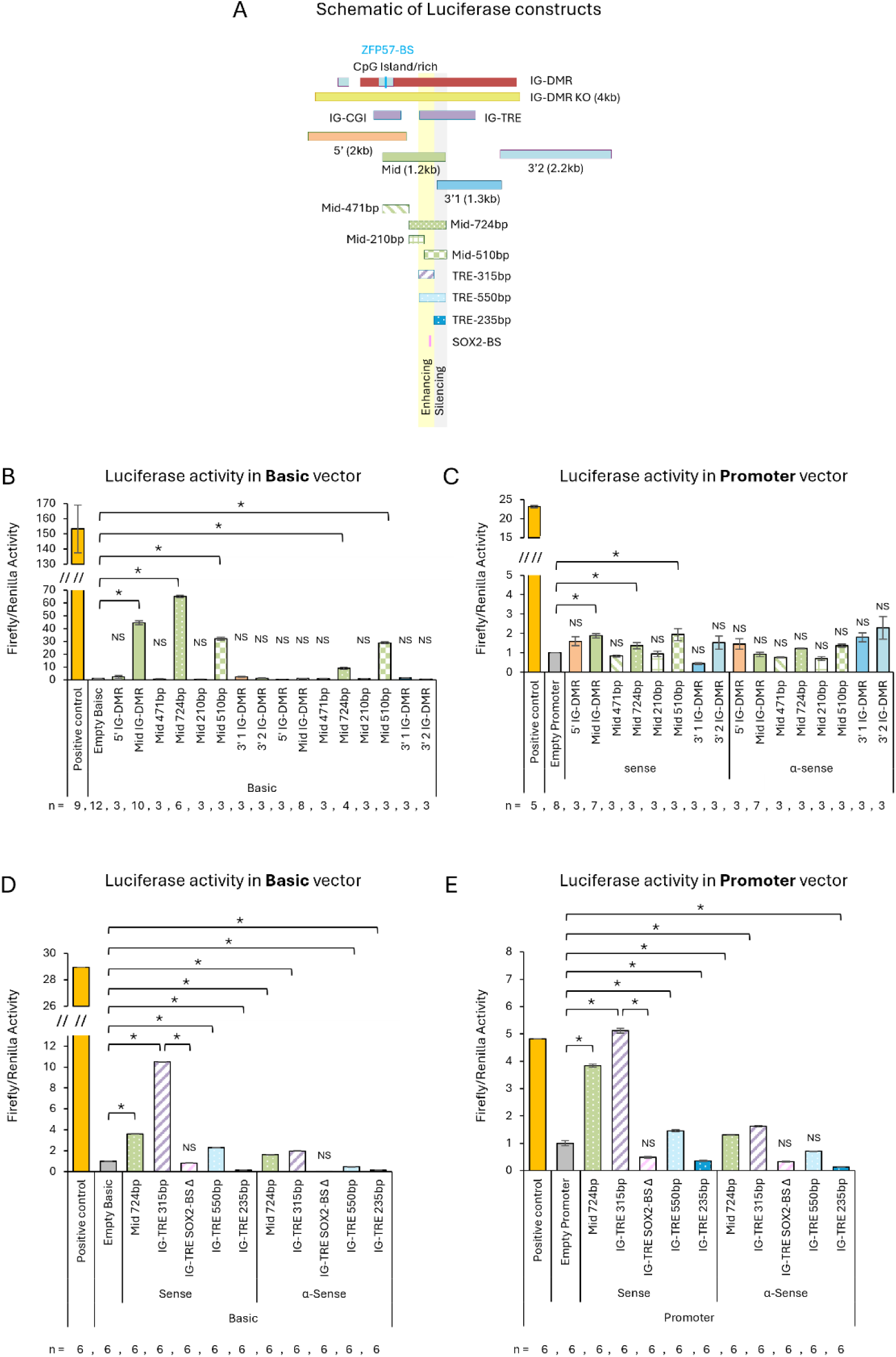
Dual activity at a minimal 550bp region at the IG-TRE: The 315bp region, and SOX2 binding within it, has enhancer like function, while the downstream 235bp region is silencing. **A**. Schematic of the IG-DMR fragments tested for activity by a luciferase reporter assay. Each fragment was cloned in both sense and antisense orientation into pGL3 vectors upstream of the firefly luciferase reporter gene. **B+C**. Graphical representation of the luciferase activity in the pGL3 Basic and Promoter vectors. The Mid 724bp fragment shows activity in basic and enhancer vectors in sense orientation only, suggesting promoter activity. **D+E**. Graphical representation of the luciferase activity in the pGL3 Basic and Promoter vectors. The Mid 315bp fragment shows activity in basic and promoter vectors, similar to the 724bp fragment, suggesting enhancer activity. This activity is lost upon deletion of the SOX2-binding site from within the 315bp fragment. In parallel, the 235bp fragment, downstream to the 315bp region, shows silencing activity, indicating dual activity in a minimal region of 550bp in the IG-TRE with both enhancing and silencing regulation. Each construct was measured twice in triplicates wells. The firefly luciferase values were normalized to the *Renilla* values and then each test construct was normalized to the corresponding empty vector. NS-not significant. Asterisks indicate statistical significance in comparison to the empty vector using a two-tailed unpaired Student’s t-test.

The 1.2kb Mid fragment (mid IG-DMR in Fig. 2) revealed promoter-like and some enhancer activity in mESCs only in the sense orientation compared to the empty vectors (Fig. 2B,C, p=0.004). Activity was not detected from any of the antisense constructs.

In order to narrow down the positive regulatory region, the Mid fragment was divided into smaller sections. The original 1.2kb Mid fragment was first separated into two fragments; 471bp (Mid-471) and 724bp (Mid-724) and assayed in ES cells. The results showed that the promoter-like activity was present in the Mid-724 (sense orientation) while the Mid-471 fragment had no activity (Fig. 2B,C and Fig. S1A, p=0.0005). Next, to further test the minimal region of activity, the 724bp Mid region was divided into two additional fragments; 210bp (Mid-210) and 510bp (Mid-510). No activity was observed from Mid-210, while promoter-like activity was seen in Mid-510. However Mid-510 had a lower level of activity than that of Mid-724 (Fig. 2B,C and Fig. S1A, p=1.08e^-08^). These results reveal that the 724bp fragment is the minimal region for effective positive regulation which the smaller fragments alone could not achieve.

### Deleting the IG-TRE in vitro results in dysregulation of the maternal epigenotype

To determine the role of the transcriptional regulatory element in regulation of expression at the Dlk1-Dio3 locus, we created mESCs deleted for the IG-TRE (Fig. 3A). We targeted 1.3kb downstream of the IG-CGI, with two guide RNAs flanking this region, as defined by Aronson *et al*^21^. Using SNPs between C57BL/6 and CAST/EiJ alleles in hybrid mESCs, we screened for clones with maternal, paternal or biallelic deletions. In addition to the full 1.3kb deletion, by chance we also obtained a smaller maternal deletion of 315bp (Fig. 3B).

**Figure 3-.**
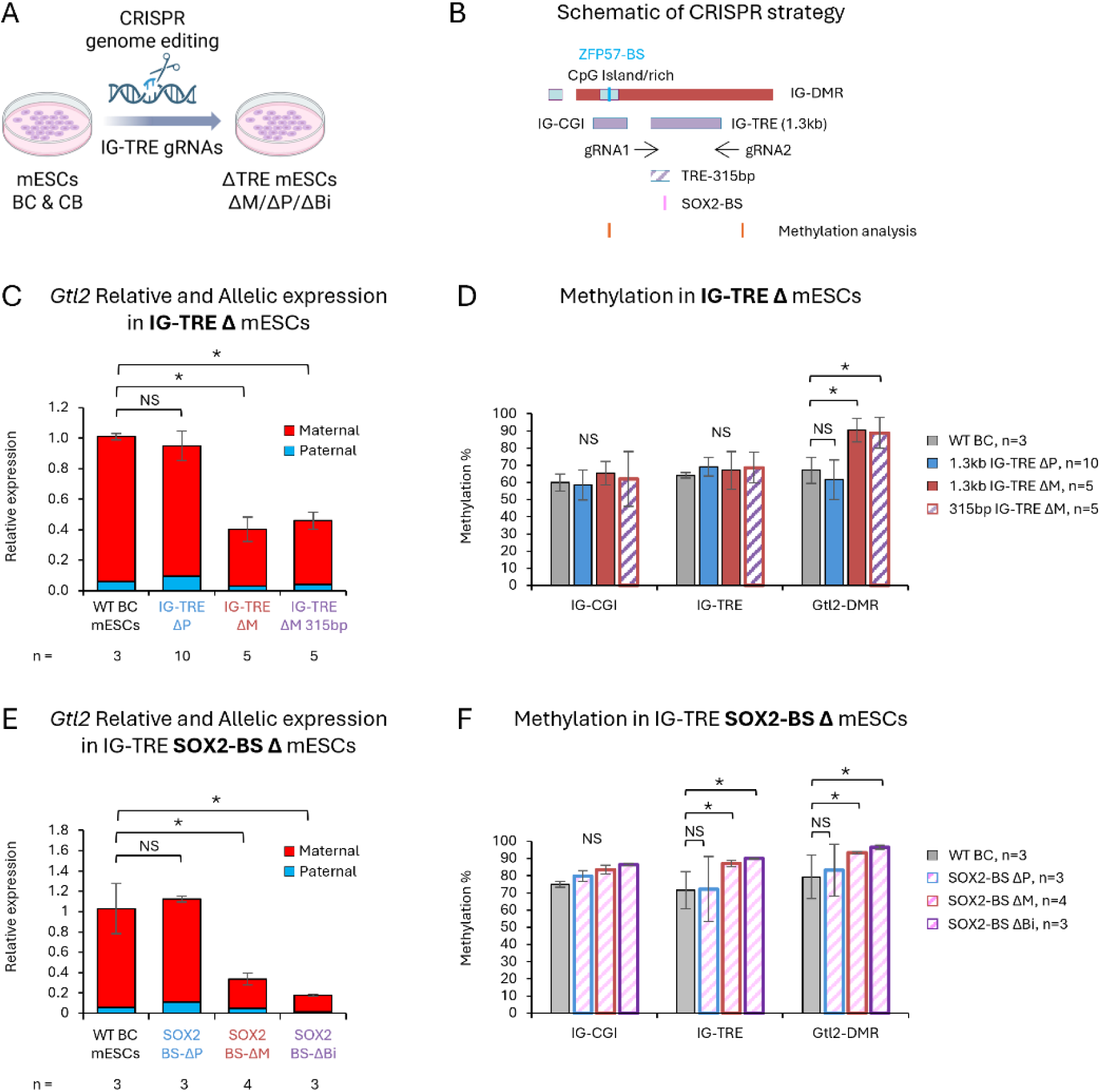
Deleting the IG-TRE *in vitro* results in dysregulation of the maternal epigenotype. **A.** Description of the CRISPR genome editing strategy in mESCs. **B**. Schematic of the gRNA locations, TRE deletions, SOX2-binding site and methylation analysis regions. **C**. Graphical representation of *Gtl2* expression in mESC clones deleted for the entire IG-TRE (1.3kb) showing major reduction in *Gtl2* expression upon maternal or biallelic deletions. **D**. Graphical representation of the methylation at three regulatory elements at the Dlk1-Dio3 domain, analyzed by bisulfited pyro sequencing, shows increased methylation at the Gtl2-DMR upon deletion of the entire 1.3kb IG-TRE and even just 315bp from the maternal chromosome. **E**. Graphical representation of *Gtl2* expression in mESC clones deleted for the SOX2-binding site shows major reduction in *Gtl2* expression upon maternal or biallelic deletions. **F**. Graphical representation of the methylation at three regulatory elements at the Dlk1-Dio3 domain, analyzed by bisulfite pyro sequencing, shows increased methylation at the Gtl2-DMR upon deletion of the SOX2-binding site from the maternal and not paternal chromosome. NS-not significant. Asterisks indicate statistical significance in comparison to wildtype using a two-tailed unpaired Student’s t-test.

Both maternal and biallelic deletions of the entire 1.3kb resulted in loss of maternal expression of *Gtl2*, *Rian* and *Mirg* (Fig. 3C, *Gtl2* p_ΔM_=0.001; p_ΔBi_=0.04 and Fig. S2A,B, *Rian* p_ΔM_=0.05; p_ΔBi_=0.05, *Mirg* p_ΔM_=0.0008; p_ΔBi_=2.4e^-05^), indicating that this region positively regulates the imprinted genes on the maternal chromosome. The same deletion on the paternal chromosome did not result in any change to gene expression at the Dlk1-Dio3 locus.

Furthermore, methylation analysis showed that loss of the maternal 1.3kb IG-TRE, by either maternal or biallelic deletions, resulted in increased methylation at the downstream Gtl2-DMR, while no change was observed in the upstream CpG island (IG-CGI) (Fig. 3D, p_ΔM_=0.005; p_ΔBi_=0.05).

Deleting only 315bp at the 5’ end of the TRE, had the same effect on maternal gene expression and methylation at the Gtl2-DMR as deleting the entire 1.3kb (Fig 3C,D) indicating that the regulatory function of the IG-TRE is controlled by elements within that minimal 315bp region.

### SOX2 binding at the IG-TRE controls maternal gene expression

To investigate which elements within the IG-TRE are responsible for maintaining the maternal epigenotype, we scanned the 315bp region for transcription factor binding sites and identified a predicted binding site for the SRY-Box Transcription Factor 2 (SOX2).

We created mESC clones with targeted deletions only for the SOX2 binding site (12bp) within the 315bp of the IG-TRE (Fig. 3B). This deletion on the maternal chromosome resulted in similar dysregulation of the maternal epigenotype, with decreased maternal gene expression (Fig. 3E, *Gtl2* p_ΔM_=0.003; p_ΔBi_=0.004 and Fig. S2C,D, *Rian* p_ΔM_=0.0005; p_ΔBi_=0.0007, *Mirg* p_ΔM_=0.006; p_ΔBi_=0.01) and increased methylation at the IG-TRE and Gtl2-DMR (Fig. 3F, IG-TRE p_ΔM_=0.009; p_ΔBi_=0.01; Gtl2-DMR p_ΔM_=0.02; p_ΔBi_=0.02). No effect was observed with deletion on the paternal chromosome. This indicates that SOX2 binding plays a key role in the cis-regulation of the maternally expressed genes in mESCs.

### The minimal 315bp region of the IG-TRE, and SOX2 binding within it, has enhancer like function, while adjacent to it there is opposing silencing function

We dissected the previously identified 724bp fragment into the 315bp fragment, identified by our CRISPR deletion experiments, and 235bp which includes the rest of the downstream sequence, as well as a 550bp fragment which encompasses both the 315bp and 235bp regions (Fig. 2A) and conducted luciferase assays *in vitro* to validate the ES cells findings.

Similar to the 724bp fragment, the minimal 315bp fragment exhibited significant activity in the sense direction, indicating promoter-like function, as well as enhancer-like function in the promoter vector (Fig. 2D,E and Fig. S1B,C, p_(Basic)_=1.16E^-05^; p_(Promoter)_=0.0001; p_(Enhancer)_=0.007). However, when the SOX2-binding site was deleted from this minimal 315bp fragment, this activity was lost (Fig. 2D,E, p_(Basic)_=0.001; p_(Promoter)_=0.0003; p_(Enhancer)_=0.02).

Strikingly, the 235bp fragment, located downstream of the 315bp region, exhibited the opposite silencing activity (Fig. 2D,E and Fig. S1B,C, p_(Basic)_=0.001; p_(Promoter)_=0.002; p_(Enhancer)_=0.0007). And finally, the 550bp fragment exhibited a partially enhancing function, reflecting a combination of the enhancing elements from the 315bp region and silencing from the 235bp region (Fig. 2D,E and Fig. S1B,C, p_(Basic)_=0.0008; p_(Promoter)_=0.01; p_(Enhancer)_=0.001).

We concluded that the 315bp region at the IG-TRE is functionally important for expression of the imprinted non-coding RNAs from the maternal chromosome, and that this function is mediated by SOX2 binding. In contrast, the 235bp region downstream of the 315bp region contains a silencing element, and the combined 550bp region is the minimal region of activity containing both regulatory elements at the IG-TRE.

### Maternal binding of ZFP281 together with SOX2 binding regulates expression on the maternal chromosome

In order to verify SOX2 binding at this predicted site, and to determine whether it binds to one or both chromosomes, we performed chromatin-immunoprecipitation and quantitative polymerase chain reactions (ChIP-qPCR) for SOX2 in wildtype and SOX2-BS mutant hybrid mESCs (Fig. 4A). Surprisingly, we found that in ESCs, SOX2 binds to both the maternal and paternal copies of the IG-TRE at the 315bp region with the maternal SOX2 binding site deletion resulting in loss of binding on the maternal copy (Fig. 4B, p=0.0007 and Fig. 4C, p=0.05). SOX2 binding at *Ceacam1* and *Scmh1* were assayed as negative and positive controls, respectively (Fig. 4B).

**Figure 4-.**
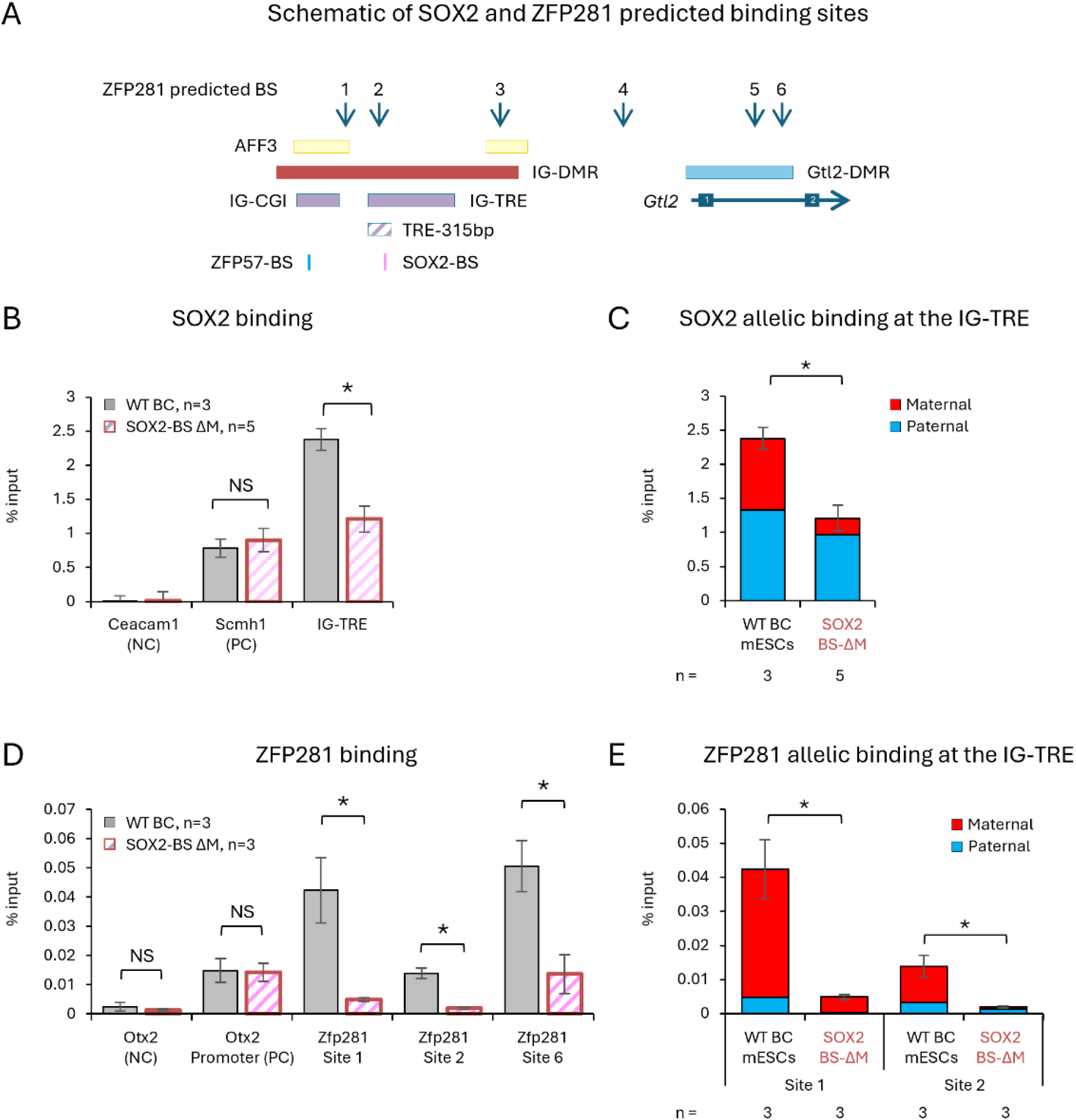
SOX2 binding together with ZFP281 regulates expression on the maternal chromosome. **A**. Schematic of SOX2 and ZFP281 predicted binding sites across the IG-DMR and Gtl2-DMR, assayed by ChIP-qPCR in B-E. **B**. Graphical representation of SOX2 binding at the IG-TRE, determined by chip-qPCR, shows depletion of SOX2 binding when the SOX2-binding site is deleted from within the 315bp region. **C**. Graphical representation of SOX2 allelic expression at the IG-TRE, as measured by ChIP-qPCR and pyro SNP analysis, shows biallelic binding in wildtype hybrid mESCs, while the maternal portion is depleted upon deletion of the maternal SOX2-binding site. **D**. Graphical representation of ZFP281 binding at the IG-TRE, determined by ChIP-qPCR, shows depletion of ZFP281 binding when the SOX2-binding site is deleted from within the 315bp region. **E**. Graphical representation of ZFP281 allelic binding at the IG-TRE, as measured by ChIP-qPCR and pyro SNP analysis, shows maternal specific binding in wildtype hybrid mESCs, which is depleted upon deletion of the maternal SOX2-binding site. NS-not significant. Asterisks indicate statistical significance in comparison to wildtype using a two-tailed unpaired Student’s t-test.

Since we found that SOX2 bound both parental chromosomes, we hypothesized that the maternal activity of the IG-TRE was combined with another factor acting allele-specifically. We identified a predicted binding site for ZFP281 within the minimal 315bp region (predicted binding site #2 ,Fig. 4A). Additionally, ZFP281 has binding sites at the 3’ end of the IG-CGI, upstream of the IG-TRE (predicted binding site #1), downstream of the IG-TRE where it has previously been shown to bind and recruit AFF3 (predicted binding site #3)^33^, downstream to the IG-DMR (predicted binding site #4) and at the Gtl2-DMR (predicted binding sites #5,6), which we hypothesize would all bind only on the unmethylated maternal chromosome to regulate allele-specific expression of the *Gtl2* polycistron^33^ (Fig. 4A).

We assayed ZFP281 binding at these predicted binding sites and at two control regions in the *Otx2* gene (negative and positive controls). We found no binding at sites 3,4,5 (Fig. S3), but that ZFP281 did bind at sites 1 and 2, upstream and in the 315bp region, and at site 6 in the Gtl2-DMR (Fig. 4D,E, p_(site1)_=0.02; p_(site2)_=0.003; p_(site6)_=0.02). Furthermore, we observed that this binding of ZFP281 to sites 1 and 2 was maternal specific, and that it was lost upon deletion of the SOX2-binding site at the 315bp region (Fig. 4D,E, p_(site1)_=0.02; p_(site2)_=0.03), indicating that ZFP281 binding is dependent on SOX2 binding, and together they regulate expression of the genes on the maternal chromosome.

### Generation of IG-TRE knockout mice

To elucidate the functional roles of the SOX2 and ZFP281 binding at the IG-TRE *in vivo* during mouse development, we created knockout mice deleted for 550bp of the IG-TRE, containing both functional elements that we identified *in vitro*: the minimal 315bp enhancing region, and the 235bp silencing region. Crossing IG-TRE^Δ550^ F1 deletion animals with C57BL/6 or CAST/EiJ wildtype partners allowed us to determine parent-of-origin effects of this deletion at different developmental stages in a highly controlled and reproducible manner (Fig. 5A).

**Figure 5-.**
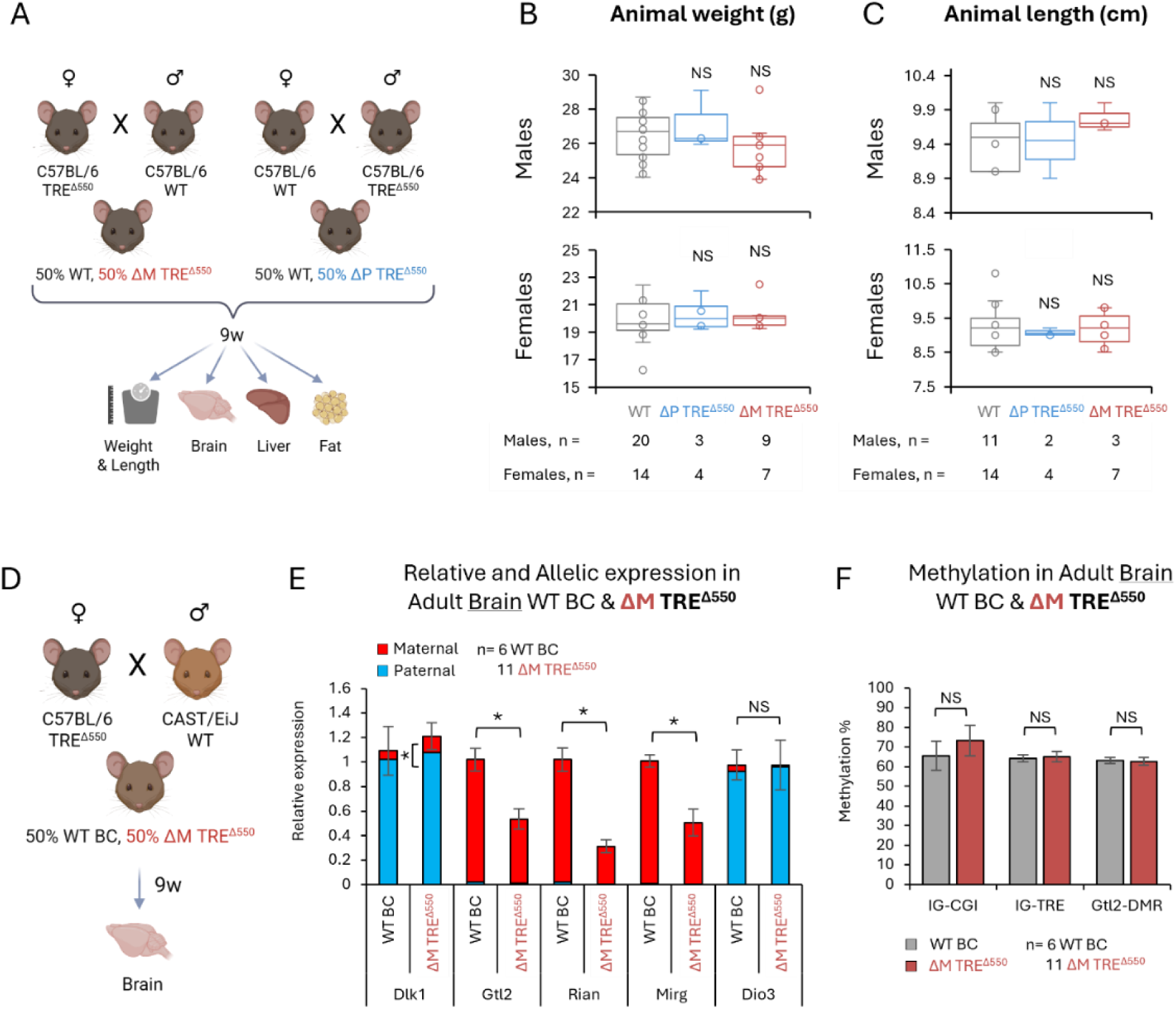
Deletion of the IG-TRE^Δ550^ *in vivo* is viable and results in decreased maternal gene expression in adult brains. **A.** Breeding strategy of the IG-TRE 550bp deletion animals crossed with C57BL/6 partners to produce maternal or paternal inheritance of the IG-TRE^Δ550^ deletion. In each litter, 50% of the pups are wildtype littermates (pure C57BL/6) and kept as controls. Tissues were collected for expression at 9 weeks. **B.** Box plot representation of total body weight (g) of 9 week-old adult animals. N_WT_=20, N_ΔP-TRE_^Δ550^=3, N_ΔM-TRE_^Δ550^=9 biologically independent males were measured; and N_WT_=14, N_ΔP-TRE_^Δ550^=4, N_ΔM-TRE_^Δ550^=7 biologically independent females were measured. **C.** Box plot representation of body length (cm) of 9 week-old adult animals. N_WT_=11, N_ΔP-TRE_^Δ550^=2, N_ΔM-TRE_^Δ550^=3 biologically independent males were measured; and N_WT_=14, N_ΔP-TRE_^Δ550^=4, N_ΔM-TRE_^Δ550^=7 biologically independent females were measured. **D.** Breeding strategy of IG-TRE^Δ550^ females crossed with CAST/EiJ males to produce hybrid animals with maternal inheritance of the IG-TRE^Δ550^ deletion. In each litter, 50% of the pups are wildtype littermates (hybrid BxC) and kept as controls. Tissues were collected for expression at 9 weeks. **E.** Graphical representation of the relative allelic expression of the Dlk1-Dio3 genes, as measured by qPCR and pyro SNP analysis, in adult brains of maternal IG-TRE^Δ550^ and BxC wildtype littermates. Expression analysis shows that *Dlk1* maternal expression is slightly increased and *Gtl2, Rian* and *Mirg* are decreased in maternal IG-TRE^Δ550^ adult brains. **F.** Graphical representation of the methylation at three regulatory elements at the Dlk1-Dio3 domain, as measured by bisulfite pyro sequencing, in embryos of all genotype groups. No change in methylation upon deletion of the IG-TRE^Δ550^ is observed. N_WTBC_=6, N_ΔM-TRE_^Δ550^=11 biologically independent adult brains were analyzed for gene expression; and N_WTBC_=6, N_ΔM-TRE_^Δ550^=11 biologically independent adult brains were analyzed for methylation. NS-not significant. Asterisks indicate statistical significance in comparison to wildtype using a two-tailed unpaired Student’s t-test.

### Phenotype and imprinting effect of the IG-TRE 550bp deletion in adult mice

Consistent with recent results^34^, we found that both maternal and paternal inheritance of the IG-TRE^Δ550^ deletion produced viable and fertile offspring. At 9 weeks of age, the weights and lengths of both maternal and paternal deleted animals were indistinguishable from their wildtype littermates (Fig. 5B,C). No difference was observed in brain and fat weight either (Fig. S4A,C,D,F), however, maternal IG-TRE^Δ550^ animals had reduced liver weight (Fig. S4B,E, p_(ΔM males)_=0.03; p_(ΔM females)_=0.04 and Fig. 6I, p_(ΔP males)_=0.03; p_(ΔM males)_=0.04; p_(ΔM females)_=0.02).

**Figure 6-.**
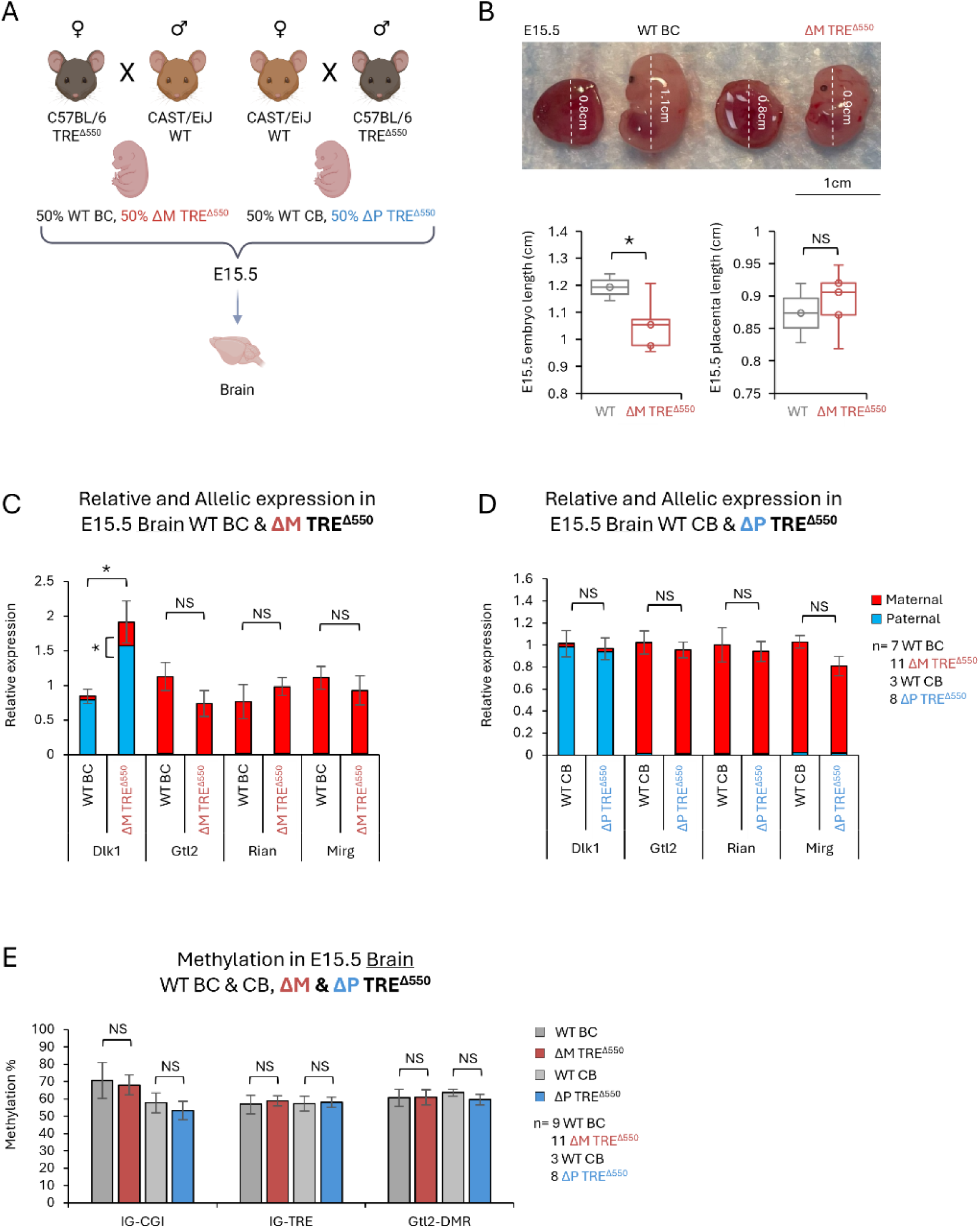
Deletion of the maternal IG-TRE^Δ550^ *in vivo* results in smaller embryos at E15.5, with increased *Dlk1* expression in the brain but no change in maternal gene expression. **A.** _Breeding strategy of IG-TRE_^Δ550^ animals crossed with CAST/EiJ partners to produce hybrid pups with maternal or paternal inheritance of the IG-TRE^Δ550^ deletion. In each litter, 50% of the pups are wildtype littermates (hybrid BxC or CxB) and kept as controls. Tissues were collected for expression at E15.5. **B.** Representative images of E15.5 embryos, embryo lengths and placentas for different genotypes. Scale bar = 1cm. **C.** Graphical representation of the relative allelic expression of the Dlk1-Dio3 genes, as measured by qPCR and pyro SNP analysis, in E15.5 brains of maternal IG-TRE^Δ550^ and BxC wildtype littermates. Expression analysis shows that *Dlk1* maternal expression is increased in maternal IG-TRE^Δ550^ E15.5 brains. Other gene expression is unchanged. **D.** Graphical representation of the relative allelic expression of the Dlk1-Dio3 genes, as measured by qPCR and pyro SNP analysis, in E15.5 brains of paternal IG-TRE^Δ550^ and CxB wildtype littermates. No change in gene expression is observed in paternal IG-TRE^Δ550^ E15.5 brains. **E.** Graphical representation of the methylation at three regulatory elements at the Dlk1-Dio3 domain, as measured by bisulfite pyro sequencing, in embryos of all genotype groups. No change in methylation upon deletion of the IG-TRE^Δ550^ is observed. N_WTBC_=7, N_ΔM-TRE_^Δ550^=11, N_WTCB_=3 and N_ΔP-TRE_^Δ550^=8 biologically independent E15.5 brains were analyzed for gene expression; and N_WTBC_=9, N_ΔM-TRE_^Δ550^=11, N_WTCB_=3 and N_ΔP-TRE_^Δ550^=8 biologically independent E15.5 brains were analyzed for methylation. NS- not significant. Asterisks indicate statistical significance in comparison to wildtype using a two-tailed unpaired Student’s t-test.

We performed allele-specific expression analysis on brains from 9 week-old hybrid animals harbouring the maternal IG-TRE^Δ550^ and their wildtype littermates (Fig. 5D). Expression of *Dlk1* from the maternally inherited chromosome, though low, was significantly increased when the IG-TRE^Δ550^ deletion was on the maternally inherited chromosome compared to wildtype where *Dlk1* is predominantly expressed from the paternal chromosome (Fig. 5E, p=0.0007). Reciprocally, *Gtl2*, *Rian*, and *Mirg* were downregulated when the deletion was on the maternal chromosome compared to controls, similar to our observations *in vitro* (Fig. 5E, p_(_*Gtl2*_)_=0.02; p_(_*Rian*_)_=0.001; p_(_*Mirg*_)_=0.001). We next analyzed methylation levels associated with the three regulatory elements at the Dlk1-Dio3 region, the IG-CGI, IG-TRE and Gtl2-DMR, and found that they maintained normal levels of methylation in maternal IG-TRE^Δ550^ adult brains compared to their wildtype littermates (Fig. 5F), indicating that the 550bp on the maternally inherited IG-TRE is involved in regulating expression from the maternal chromosome *in vivo*, but that its deletion neither changes the imprint at the ICR nor is detrimental for embryonic development.

### Tissue specific dissection of IG-TRE^Δ550^ during embryonic development

To further elucidate the effect of the IG-TRE^Δ550^ deletion in a time-dependant manner, we crossed IG-TRE^Δ550^ heterozygote animals with CAST/EiJ partners to dissect mutant embryos at E15.5 and analyzed gene expression and methylation in E15.5 embryonic brains upon maternal and paternal transmission of the deletion (Fig. 6A). We found that maternal IG-TRE^Δ550^ E15.5 embryos were smaller than their wildtype littermates, while their placenta length was unchanged (Fig. 6B, p=0.05). Similar to our observations from adult brains, we found that *Dlk1* expression was increased in maternal IG-TRE^Δ550^ E15.5 brains compared to controls (Fig. 7C, p=0.01) and that strikingly this expression came from both chromosomes. Furthermore, surprisingly and unlike adult brains and mESCs, there was no change in expression of the maternally expressed genes. No expression change of any gene at the Dlk1-Dio3 domain was observed with paternal IG-TRE^Δ550^ E15.5 brains compared to hybrid wildtype littermates (Fig. 6D). Lastly, and consistent with the adult findings, we found no change in methylation at the IG-CGI, IG-TRE or Gtl2-DMRs in E15.5 brains with either maternal or paternal deletion of the IG-TRE^Δ550^ (Fig. 6E).

**Figure 7-.**
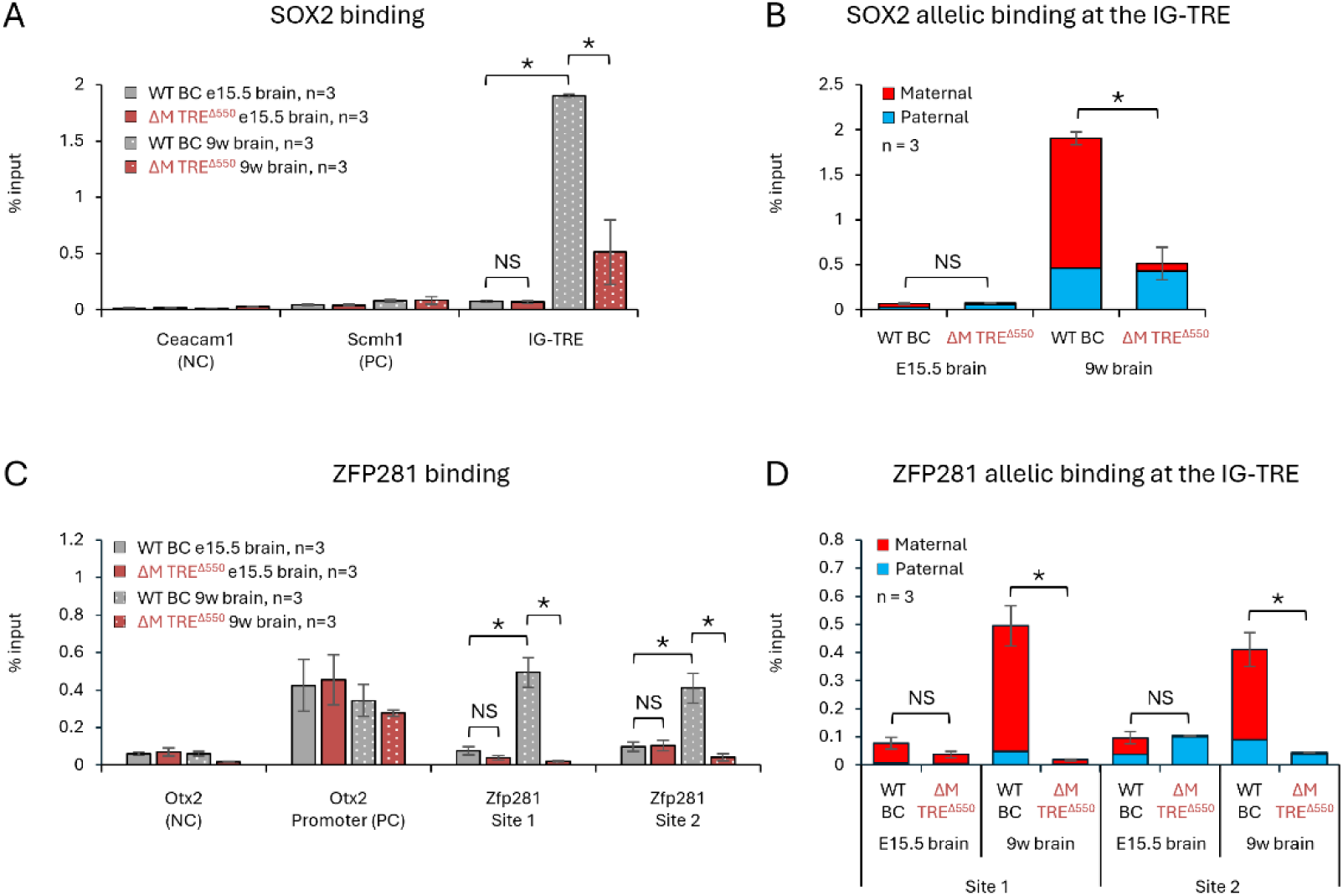
SOX2 and ZFP281 bind the maternal IG-TRE in adult brains *in vivo*. **A**. Column graph representation of SOX2 binding at the IG-TRE in embryonic and adult brain, determined by chip-qPCR, shows more binding in adult vs embryonic brains, and that SOX2 binding is reduced when the maternal IG-TRE^Δ550^, including the SOX2-binding site, is deleted. **B**. Column graph representation of SOX2 allelic expression at the IG-TRE, as measured by ChIP-qPCR and pyro SNP analysis, shows primarily maternal binding in wildtype hybrid brains, and depletion of the maternal portion upon deletion of the maternal IG-TRE^Δ550^, which includes the SOX2-binding site. **C**. Column graph representation of ZFP281 binding at the IG-TRE in embryonic and adult brain, determined by ChIP-qPCR, shows increased binding in adult vs embryonic brains, and that ZFP281 binding is depleted when the maternal IG-TRE^Δ550^, including the SOX2-binding site, is deleted. **D**. Column graph representation of ZFP281 allelic expression at the IG-TRE, as measured by ChIP-qPCR and pyro SNP analysis, shows maternal specific binding in wildtype hybrid brains, which is depleted upon deletion of the maternal IG-TRE^Δ550^, including the SOX2-binding site. NS-not significant. Asterisks indicate statistical significance in comparison to wildtype using a two-tailed unpaired Student’s t-test.

Taken together, these results indicate that the IG-TRE functions in a developmental stage-specific manner that may associate with a requirement for exquisite *Dlk1* dosage control. For example, as opposed to its role as an enhancer of maternal expression in ESCs and in adult brains where *Dlk1* is not strongly expressed, *in vivo* during embryonic development where *Dlk1* expression is high in the brain, the IG-TRE has a more pronounced silencing effect on *Dlk1* expression, and its absence results in changes to *Dlk1* expression from both chromosomes.

### SOX2 and ZFP281 bind to the maternal IG-TRE in adult brains in vivo

To explain the observed differences between the functionality of our IG-TRE^Δ550^ deletion in embryonic and adult brains, we assayed the binding of SOX2 and ZFP281 *in vivo*. We found that both SOX2 and ZFP281 bind to the IG-TRE in 9 week-old brains, but that binding is weak in embryonic brains (Fig. 7A,C, p_(SOX2)_=0.0001, p_(ZFP281 site1)_=0.0004, p_(ZFP281 site2)_=0.003). Furthermore, as opposed to the biallelic binding of SOX2 that we observed in ESCs, *in vivo* both SOX2 and ZFP281 binding was maternal specific, and binding of both was lost in IG-TRE^Δ550^ maternal deletion brains (Fig. 7B,D, p_(SOX2)_=0.0008, p_(ZFP281 site1)_=0.0001, p_(ZFP281 site2)_=0.001).

We conclude that *in vivo*, SOX2 binding at the IG-TRE is required for ZFP281 binding and maternal gene regulation in adult brains, but that this is either not required or not established yet in embryonic brains. While the maternal silencing of *Dlk1* is already functional *in utero* it is probably mediated by other elements within the 235bp of the IG-TRE.

### Balancing Dlk1 gene dosage by combining two IG-DMR mouse deletion models

In light of our findings that the IG-TRE 550bp deletion from the maternal chromosome results in increased *Dlk1* expression, but that the animals have no apparent phenotype, we asked whether this increased expression was significant enough to rescue a postnatal phenotype that arises from loss of *Dlk1* expression on the opposite chromosome. Previously we described that deleting the IG-CGI from the paternal chromosome results in an epigenotype switch from paternal to maternal like behaviour^28^, with downregulation of *Dlk1* and upregulation of the maternal transcripts *Gtl2*, *Rian* and *Mirg* (Fig. 8A). Furthermore we showed that balancing the gene dosage, through flipping expression of paternally expressed genes from the maternal chromosome and maternally expressed genes from the paternal chromosome, was sufficient for normal development and viability. However, it remains unknown whether this is due to a balanced dosage of a specific gene or a combination of multiple/all genes at the Dlk1-Dio3 domain. The new IG-TRE^Δ550^ model described here offers a rare opportunity to address the possibility that the paternal or maternal genes at the Dlk1-Dio3 domain are not equally essential for viability. Specifically, crossing IG-TRE^Δ550^ heterozygote females (increased *Dlk1*) with IG-CGI^Δ^ heterozygote males (loss of *Dlk1*) is expected to produce double deletion embryos with a balance of *Dlk1* expression, while the maternal genes will remain dysregulated (Fig. 8A). We hypothesized that inheritance of both deletions, would rescue the postnatal lethality of the paternal IG-CGI deletion if balancing *Dlk1* expression was sufficient for development, but not if balance of the maternal transcripts was essential.

**Figure 8-.**
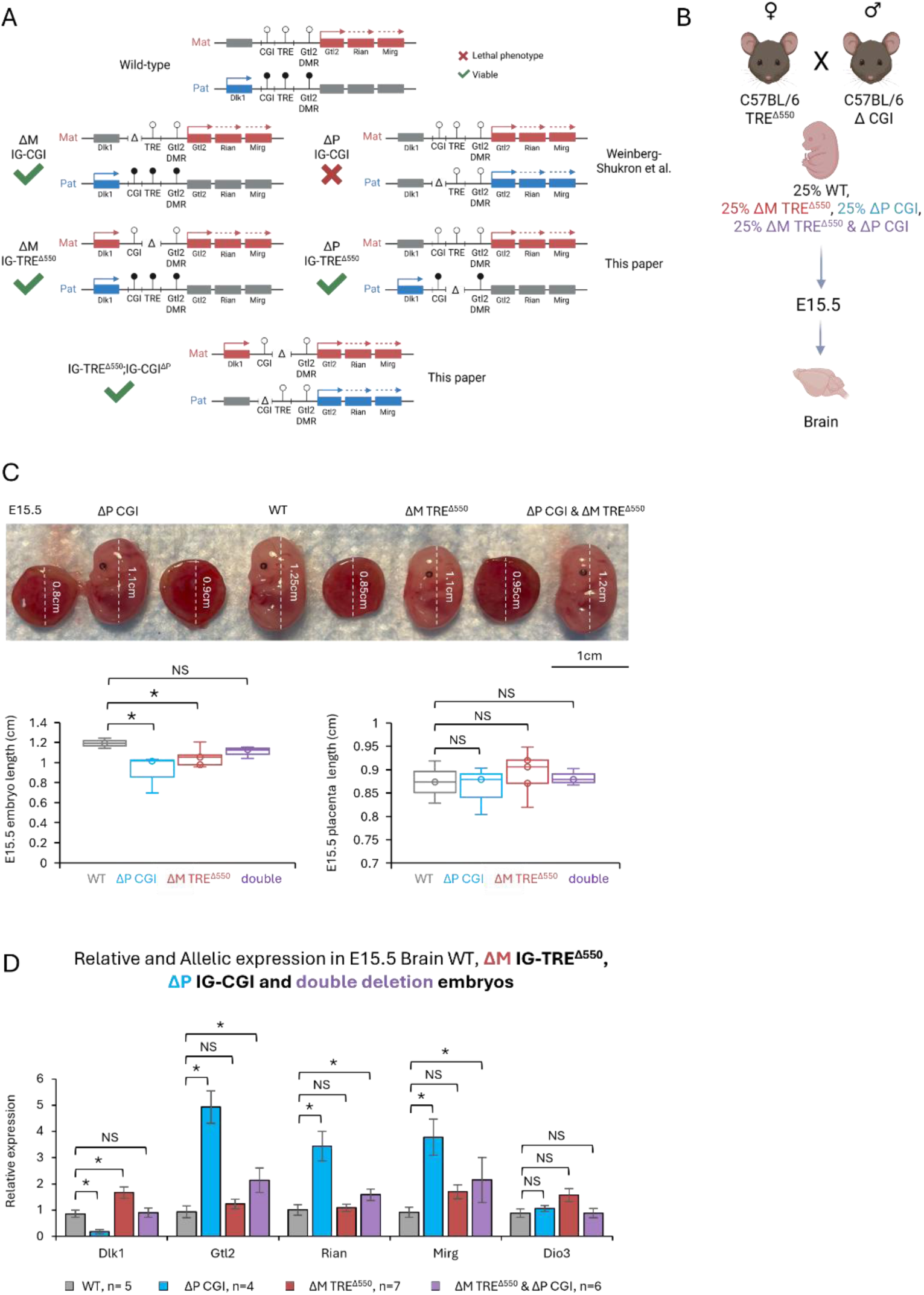
Deletion of the maternal IG-TRE^Δ550^ rescues the lethality of the paternal IG-CGI deletion *in vivo* and results in balanced gene dosage at the Dlk1-Dio3 domain. A. Schematic representation of the predicted outcome of crossing IG-CGI^ΔP^ and IG-TRE^Δ550^ mouse strains. The increased *Dlk1* expression caused by the maternal IG-TRE^Δ550^ deletion balances the loss of *Dlk1* expression caused by the IG-CGI paternal deletion and results in viable double deletion pups. **B.** Breeding strategy of IG-TRE^Δ550^ heterozygote females with IG-CGI^Δ^ heterozygote males to produce double deletion pups. In each litter, 25% of the pups are wildtype littermates (pure C57BL/6) and kept as controls. 25% of the pups are IG-CGI^ΔP^ which is lethal postnatally (as described in *Weinberg-Shukron et al*. 2022). 25% of the pups are IG-TRE^Δ550^ and 25% have both IG-CGI^ΔP^ and IG-TRE^Δ550^. Tissues were collected for expression at E15.5. **C.** Representative images of E15.5 embryos, placentas and lengths. Scale bar = 1cm. While single deletion embryos are slightly shorted in length, double deletion embryos are indistinguishable from wildtype. **D.** Graphical representation of the relative allelic expression of the Dlk1-Dio3 genes, as measured by qPCR. Gene expression in double deletion E15.5 brains is not different from wildtype. IG-CGI^ΔP^ embryos show a paternal-to-maternal epigenotype switch, as previously described, with a downregulation of *Dlk1* and upregulation of the maternal transcripts *Gtl2, Rian* and *Mirg*. Maternal IG-TRE^Δ550^ embryos show an upregulation of *Dlk1* and no change in other genes. N_WT_=5, N_ΔP-CGI_=4, N_ΔM-TRE_^Δ550^=7 and N_ΔP-CGI&ΔM-TRE_^Δ550^=6 biologically independent E15.5 brains were analyzed for gene expression; NS-not significant. Asterisks indicate statistical significance in comparison to wildtype using a two-tailed unpaired Student’s t-test.

### Viability of IG-TRE^Δ550^ and IG-CGI^ΔP^ double deletion pups

Interestingly, we found that while isolated IG-CGI^ΔP^ pups died postnatally as expected, double deletion pups were viable and overall indistinguishable from their wildtype and maternal IG-TRE^Δ550^ littermates (Fig. S5A-I). Notably, IG-TRE^Δ550^;IG-CGI^ΔP^ double deletion females had slightly reduced brain weights (Fig. S5C, p=0.01) and about 50% more fat (Fig. S5E, p=0.04) than wildtype females at 9 weeks of age, but this was not statistically significant when normalized to body weight (Fig. S5G,I).

### Imprinting analysis of IG-TRE^Δ550^;IG-CGI^ΔP^ double deletion embryos

To understand the regulatory configuration that allows these animals to develop normally, we dissected embryos at E15.5 and analyzed gene expression in embryonic brains (Fig. 8B). We found that, similar to our previous observations for isolated paternal IG-CGI^ΔP^ and maternal IG-TRE^Δ550^ embryos, mutant E15.5 embryos were smaller than their wildtype littermates, while their placenta length was unchanged (Fig. 8C, p_(ΔP CGI)_=0.05; p_(ΔM TRE)_=0.05). IG-TRE^Δ550^;IG-CGI^ΔP^ double deletion embryos however, were not significantly different to their wildtype littermates. Gene expression analysis confirmed the previously reported gene dosage effects of the isolated paternal IG-CGI^ΔP^ and maternal IG-TRE^Δ550^ deletions, with the former exhibiting lower levels of *Dlk1* relative to wildtype littermates together with elevated levels of all maternal transcripts at this locus (Fig. 8D, p_(_*Dlk1*_)_=0.005; p_(_*Gtl2*_)_=0.0008; p_(_*Rian*_)_=0.006; p_(_*Mirg*_)_=0.007), while the latter shows increased *Dlk1* expression and no change to the maternal genes (Fig. 8D, p_(_*Dlk1*_)_=0.03). Importantly, we found that *Dlk1* expression is restored to normal levels in IG-TRE^Δ550^; IG-CGI^ΔP^ double deletion embryos, while maternal gene expression remains elevated (Fig. 8D, p_(_*Gtl2*_)_=0.05; p_(_*Rian*_)_=0.05; p_(_*Mirg*_)_=0.05).

Combining the results of all the ICR deletion models, shows that loss of both the paternal gene *Dlk1*, as in the IG-CGI^ΔP^ model, or the maternal genes *Gtl2*, *Rian* and *Mirg*, as in the IG-DMR^ΔM^ model, do not allow for normal development. Increased *Dlk1* expression, however, as in the maternal IG-TRE^Δ550^ model, has no apparent mutant phenotype, suggesting that overexpression is not as detrimental as loss of expression. Moreover, reintroducing *Dlk1* expression by combining the maternal IG-TRE^Δ550^ and the IG-CGI^ΔP^ deletions, elevates *Dlk1* sufficiently to rescue the lethality of the IG-CGI^ΔP^ alone.

Our results, therefore highlight a fundamental requirement for expression of the Dlk1-Dio3 genes from at least one chromosome, regardless of parental origin, for proper embryonic growth and survival and that the lethality observed in the IG-CGI^ΔP^ and IG-DMR^ΔM^ deletion models is due to loss of expression *of Dlk1* and not overexpression of the opposite maternally expressed non-coding RNA genes.

## Discussion

Appropriate allele specific expression of the genes at the Dlk1-Dio3 region is necessary for normal mammalian development. Understanding the regulation of genomic imprinting at the Dlk1-Dio3 domain requires *in vivo* models that faithfully recapitulate developmental complexity. Assessing different models that remove various elements in the region, has allowed us to dissect the chain of events that is necessary to establish and maintain imprinted gene expression.

While a full deletion of the IG-DMR resulted in a maternal-to-paternal switch, an isolated deletion of the IG-CGI alone resulted in the reciprocal paternal-to-maternal switch. The remaining part of the IG-DMR, the IG-TRE, has been suggested to control maternal expression, but this was not demonstrated *in vivo* until recently^34^. Matsuzaki *et al*. showed that a mouse IG-DMR transgene became highly methylated from both copies, suggesting that the IG-DMR fragment tested did not protect the maternal IG-DMR from genome-wide *de novo* DNA methylation^40^. Interestingly, the fragment did not contain most of the IG-TRE, indicating that one function of the IG-TRE may be to protect the maternal sequence from global *de novo* methylation after implantation.

Here, we defined the minimal region of enhancer activity required for maternal epigenotype maintenance to 315bp at the 5’ of the IG-TRE and found that this activity is dependent on binding of SOX2 and ZFP281. Maintenance of hypomethylation at the maternal *H19*-ICR is also known to involve SOX/OCT factors^41^. We show that the function of the maternal IG-TRE is similarly mediated by SOX2 binding and when the SOX2 binding site was deleted, the maternal IG-TRE could not bind ZFP281, leading to decreased expression of the maternal gene *Gtl2*. We showed that these elements also bind to the IG-TRE *in vivo* with increased binding in adult brain compared to embryonic brain.

However, our IG-TRE^Δ550^ deletion which includes the 315bp region resulted in no overt phenotype in mice, highlighting a divergence between *in vitro* and *in vivo* imprinting control. Our *in vitro* assays also identified a silencing element within the IG-TRE, but since *Dlk1* is not expressed in ESCs we could not test the complete effects of this region in culture, further emphasizing the importance of studying these elements *in vivo*. While embryonic stem cell-based assays have provided insights into transcription factor binding and epigenetic regulation, our data demonstrate that *in vitro* systems do not fully capture the functional consequences of regulatory element deletions *in vivo*. Similar discrepancies were previously reported for AFF3/ZFP281 activity, where this binding regulated maternal expression in ESCs but not *in vivo*^25,33^, suggesting that the effect of ZFP281 binding at the IG-DMR is temporal and cell-type specific. Similarly, we also observed differences in allelic binding of SOX2, where it bound to both chromosomes in ESCs, but was maternal-specific in brain tissue.

The 550bp IG-TRE deletion described here did not recapitulate the maternal lethality of the full IG-DMR deletion^18^ or the 2.7kb IG-TRE deletion^34^ *in vivo*, enabling refinement of the functional footprint of the IG-TRE by targeting a smaller element than in previous studies (Fig. 9). Our results show that deletion of the IG-TRE^Δ550^ in adult brains results in reduced maternal expression, suggesting that this region of the IG-TRE does have enhancer function on the maternal chromosome (similar to our observations *in vitro*), but that *in vivo* this is not required during embryonic development. We also found that SOX2 and ZFP281 bind at the IG-TRE *in vivo*, and that their binding in adults and not *in utero* explains the observed differences in gene expression caused by the IG-TRE^Δ550^ between these stages. Taken together, these results suggest that the IG-TRE controls dosage at the Dlk1-Dio3 region in a time-specific manner. In the embryo, when SOX2 and ZFP281 do not bind at the IG-TRE and *Dlk1* levels are higher, the silencing function modulates *Dlk1* dosage. In contrast, in the adult brain and in mESCs, when total *Dlk1* expression is low, the IG-TRE functions as an activator of *Gtl2*, mediated by SOX2 and ZFP281 binding (Fig. 9).

**Figure 9-.**
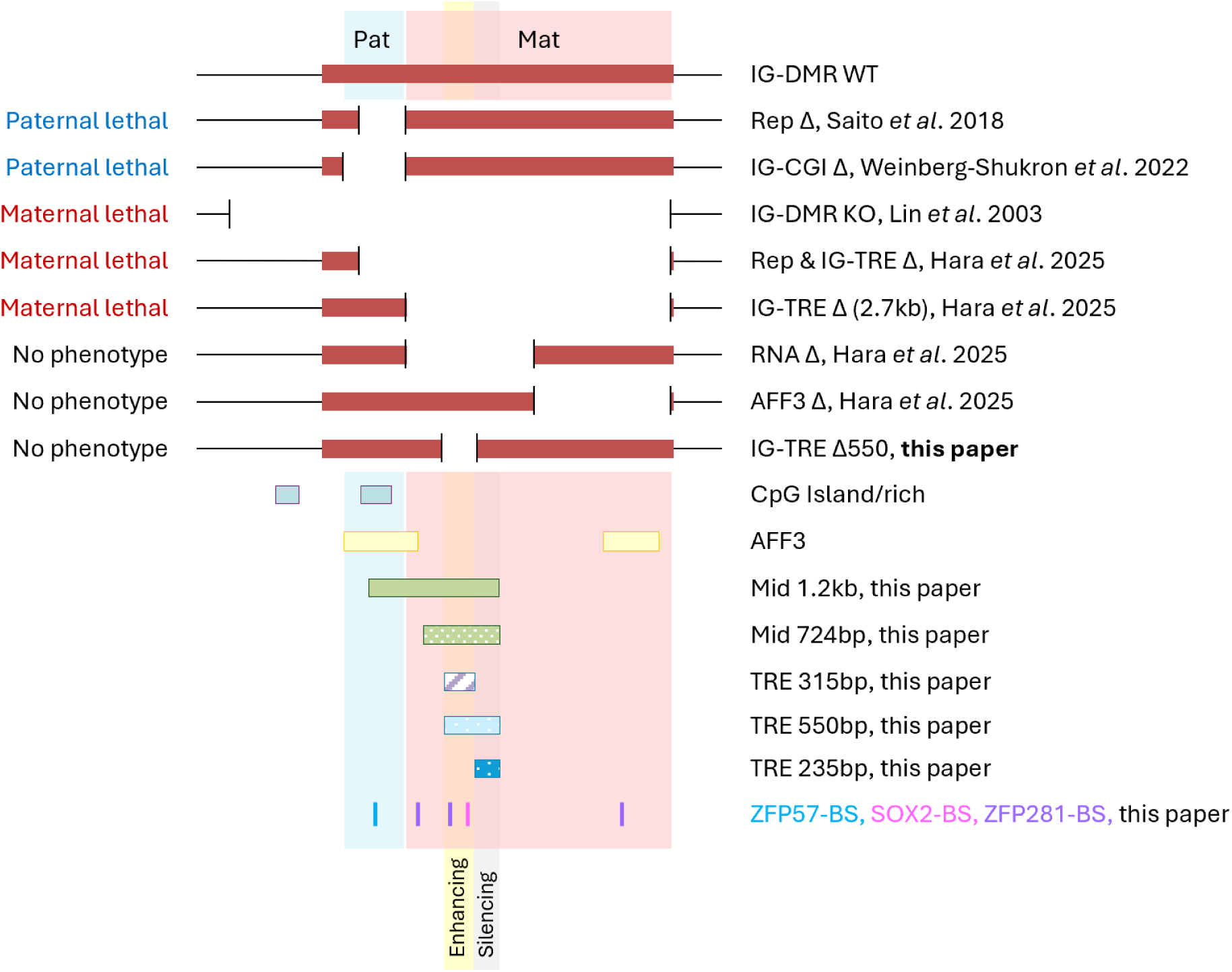
An integrated summary depicting the allele-specific counterparts involved in cis-regulation at the Dlk1-Dio3 region. Regulation of the paternal epigenotype involves the CpG island and ZFP57 binding site (BS). While maternal regulation is more complex and involves multiple transcriptional regulatory elements (TRE) downstream to the CpG island, including: AFF3, SOX2 binding and ZFP281 binding. Deletions of these elements on their own (RNA Δ model, AFF3 Δ model or the IG-TRE^Δ550^ model in this paper) have no phenotype and do not dysregulate imprinting on the maternal chromosome. The minimal region of 550bp at the IG-TRE includes dual enhancing and silencing activity, positively driving maternal gene expression and reducing paternal expression from the maternal chromosome.

Furthermore, while the 550bp IG-TRE deletion did not disrupt maternal gene expression *in utero*, it caused a significant upregulation of *Dlk1* from both chromosomes, suggesting that the silencing element in the deleted region contributes to dosage restraint *in cis* and *in trans*, albeit redundant with other elements regulating maternal gene expression, thus allowing normal development.

This finding points to a trans-allelic regulatory effect at the Dlk1-Dio3 locus, whereby deletion of a regulatory element on the maternal chromosome results in increased expression from the paternal allele. Rather than acting solely *in cis*, the IG-TRE appears to participate in inter-allelic communication, indicating that imprinting control at this locus involves coordinated regulation across both parental chromosomes. *Gtl2* RNA has been implication in this previously, however further interrogation of this is required in our models in a cell and stage-specific manner. Such cross-chromosomal influence adds an additional layer of complexity to imprinting regulation and may help explain why perturbations at imprinting control regions often produce reciprocal effects that are difficult to interpret in terms of simple *cis*-acting mechanisms alone.

Importantly, our double deletion model, which combined the maternal IG-TRE 550bp deletion with our previously characterized paternal IG-CGI deletion, provided functional evidence for regulatory compensation between maternal and paternal control elements. While the paternal IG-CGI deletion alone reduces *Dlk1* expression, increases maternal gene expression, and leads to embryonic lethality, this phenotype is fully rescued by simultaneous deletion of the maternal IG-TRE^Δ550^. These findings support a model in which loss of expression is more detrimental than gain, suggesting that precise regulation of *Dlk1* is required within a narrow threshold range, and that modest overexpression is better tolerated than deficiency.

It is important to note that *Dlk1* knockout mice are not lethal but display metabolic phenotypes as well as overall growth retardation^42^. Hence, the lethality of the IG-CGI paternal deleted pups could not be attributed to loss of *Dlk1* expression alone. The maternalization of the paternal chromosome by the IG-CGI deletion results in loss of expression of other paternal genes as well, such as *Rtl1* and *Dio3*, and their contribution to the lethality of these embryos should also be considered.

Together, this work refines the regulatory map of the Dlk1-Dio3 domain and identifies novel transcription factor interactions at the IG-TRE and both enhancing and silencing elements. It provides *in vivo* evidence for epigenetic redundancy of cis-regulatory elements at this region and underscores the necessity of *in vivo* models for deciphering the complex chain of epigenetic events that are required to establish and maintain imprinted gene expression throughout development.

## Materials and Methods

### Cloning and generation of Luciferase vectors

#### Polymerase Chain Reaction (PCR)

PCR fragments were amplified using Novagen KOD Hot Start DNA polymerase. 1µl of gDNA from C57BL/6J and CAST/EiJ mESCs hybrids was used as template in a 20µl reaction containing 1XPCR Buffer (KOD Hot Start, Novagen#71155), 200µM dNTPs (Novagen#71154), 1.8mM MgSO_4_ (Novagen#71156), 0.6mM DMSO, 5.8mM cresol red, 0.5U of Hot Start KOD polymerase (Novagen#71316) and 0.6µM of each primer. Primers introduced NheI or KpnI restriction sites at each end in order to clone both in sense and antisense orientation. The PCR program was 95°C for 2’, followed by 35-cycles at 95°C for 20’’, 60°C for 15’’ and 70°C for 20’’. The PCR products were visualized by agarose gel electrophoresis in TBE buffer and stained with SafeView (NBS Biologicals #NBS-SV). Primers used to generate the luciferase constructs are listed in Table S1.

#### A-Tailing of PCR products

KOD PCR products we A-tailed prior to cloning. Each 25µl reaction contained 12.5µl PCR product, 1XHotStarTaq master mix (Qiagen#1005479), 0.2mM dATP and 5U HotStarTaq (Qiagen#1032502). The Taq polymerase was activated at 95°C for 15’. A-tailing reaction was then performed at 70°C for 30’.

#### TA Cloning

The A-tailed PCR product was cloned into the pGEM-T Easy Vector System (Promega#A1360) using the manufacturer’s protocol, with a 1:3 vector:insert molar ratio. A 10µl reaction contained the according amount of PCR product, 50ng of vector, 1Xreaction buffer and 2U T4 ligase. For ligation the reaction was incubated at 4°C overnight. 2µl of the ligation reaction was added to a sterile 1.5ml microcentrifuge tube containing 50µl of competent E. coli cells (Stellar™ Competent Cells, Takara Bio#636763) on ice. The mix was incubated on ice for 30’ and then heat shocked for 45’’ at 42°C. After another 2’ incubation on ice, 500µl of SOC medium (Takara Bio#ST0215) was added to the tubes followed by 1h incubation at 37°C. As pGEM-T Easy Vectors allow blue/white selection, 100µl of ligation reaction was plated on 100µg/ml ampicillin (Sigma-Aldrich#A9518), 20µg/µl X-gal (MELFORD#B71800), 100µM IPTG (MELFORD#I56000) plates. These plates were incubated at 37°C overnight.

#### Cloning

PCR products were digested with appropriate restriction enzymes by adding 1Xrestriction buffer (rCutSmart™ Buffer, New England Biolabs#B6004) and 20-50U of enzyme (NheI-New England Biolabs#R3131, KpnI-New England Biolabs#3142 or SacI-New England Biolabs#R3156), then incubated at 37°C overnight followed by purification using QIAquick PCR Purification Kit (Qiagen#28106) or, in case of pGEM-T Easy Vectors, directly from an agarose gel using the MinElute Gel Extraction Kit (Qiagen#28606) according to the manufacturer’s protocol.

The fragments were cloned into three different pGL3 vectors (Promega) which contain the Firefly luciferase reporter gene. The three vectors are pGL3-Basic (containing no promoter or enhancer, Promega#E1751), pGL3-Promoter (containing a SV40 promoter element, Promega#E1761) and pGL3-Enhancer (containing a SV40 enhancer element, Promega#E1771). 4µg of these vectors were digested by NheI and KpnI restriction enzymes overnight at 37°C. Digests were checked visually by gel electrophoresis to ensure complete digestion. After complete digestion, the enzyme was heat inactivated at 65°C for 20’. 50ng of the vector was ligated with an appropriate amount of insert to reach a 1:3 vector:insert molar ratio, using T4 ligase (New England Biolabs#M0202) and incubated at 16°C overnight. Transformation was performed as described above.

#### Colony PCR

Colony PCR was performed in 0.2ml microcentrifuge tubes or 96-well plates. Each 25µl reaction contained 1XPCRBIO HS Taq Mix (PCR Biosystems#PB10.23-02), 10µM of each insert specific primer and 5µl of the water/bacteria mix as a template. The PCR program was 95°C for 10’, followed by 25 cycles of 95°C for 15’’, 60°C for 15’’, 72°C for 15’’ and a 5’ extension cycle at 72°C. The PCR products were visualized by agarose gel electrophoresis in TBE buffer and stained with SafeView.

#### Plasmid DNA minipreps

3-5ml of LB medium containing 100µg/ml ampicillin was inoculated with 200µl LB medium containing half a positive bacterial colony and was grown at 37°C with shaking at 200rpm overnight. DNA was prepared using the ZR Plasmid Mini/Midiprep-Classic Kit (Zymo Research#D4016/#D4200) according to the manufacturer’s protocol. The plasmid DNA was verified by sequencing using vector specific primers described in Table S1.

#### Transfection

Before transfection of luciferase plasmids into mESCs, MEF feeder cells were depleted on 0.2% gelatin for 40’ at room temperature. 1-2x10^4^ cells were seeded in 0.2% gelatin coated wells of a 48-well plate and incubated for 6h at 37°C. For transfection, 500ng of Firefly and 1ng of *Renilla* plasmids were incubated together with 25µl OptiMEM (Gibco#31985062) and 1µl *Trans*IT-X2® (Mirus#MIR6000) for 20’ at room temperature. After that, the reaction was added dropwise to the mESCs. Fresh media was added on the next day. After 24h cells were taken for luciferase assays. To generate comparable results, each plate contained a positive control (pGL3-Control, Promega#E1741), negative control (pRL-SV40 *Renilla* only, Promega#E2231) and an empty vector.

#### Dual-Luciferase® Reporter Assay System

The Dual-Luciferase® Reporter (DLR^TM^) Assay System (Promega#E1910) allowed for the measuring of the luminescence for Firefly and then *Renilla* sequentially with a single transfection reaction^43^. The DLR assay was performed 24h after transfection. Cells were prepared by washing twice with PBS and then lysed with 65µl/well 1XPassive Lysis buffer for 15’ on a rocking platform. To assess firefly luminescence, 10μl of lysed cells were added to 50μl Luciferase Assay Reagent (LARII) and put in a single channel TD20/20 Luminometer (Turner Designs) to measure the signal. The luminometer was programmed with a 2’’ pre-read delay, followed by a 10’’ measurement period. After Firefly, *Renilla* luminescence was read by adding 50μl of Stop&Glo® substrate, which first quenches firefly reaction and then activates *Renilla*. Each construct was measured twice in triplicates wells. The firefly luciferase values were normalized to the *Renilla* values and then each test construct was normalized to the positive control and then plotted as fold-change compared to the empty vector.

### mESCs Cell Culture

C57BL/6J x CAST/EiJ hybrid mESCs were cultured at 37°C with 5% CO2, on plates coated with 0.2% gelatine on irradiated mouse embryonic fibroblasts (iMEFs, DR4), in standard ESCs medium: 500ml DMEM (Gibco#31966-021) supplemented with 20% US certified FBS (Gibco#10270-106), 10ug recombinant leukemia inhibitory factor (LIF, Merck#ESG1107), 0.1mM beta-mercaptoethanol (Gibco#31350-010), penicillin/streptomycin 1mM (Sigma-Aldrich#P0781), L-glutamine (Gibco#25030-024) and 1% nonessential amino acids (Gibco#11140-050).

### Generation of IG-TRE deletions in mESCs

#### Cloning

To establish mESCs deleted for the IG-TRE and for the SOX2-binding site at the IG-TRE, targeting vectors and CRISPR/Cas9 plasmids were cloned and co-transfected into mESCs. gRNAs were selected for minimal off targets using CCTop-CRISPR/Cas9 target online predictor (https://crispr.cos.uni-heidelberg.de/, Stemmer 2015). sgRNA oligos were duplexed to double strands and phosphorylated by T4 PNK to allow subsequent cloning into Cas9 coding plasmids. Each 10µl reaction contained 10µM of each oligo (forward and reverse), 1mM ATP (New England Biolabs#P0756), 1Xreaction buffer and 0.5U T4 PNK (New England Biolabs#M0201). The reaction was incubated at 37°C for 30’, 95°C for 5’ and then cooled down to 25°C at 0.1°C/second. The phosphorylated DNA products were cloned into the pSpCas9(BB)-2A-Puro plasmid (Addgene#PX459), which expresses *Cas9* under the control of a U6 promoter and includes an ampicillin and puromycin resistance for following selections. Cloning was done by BbsI restriction enzyme digest and T4 ligase mediated ligation. Each 20µl reaction contained 100ng Cas9 vector, 2µl of 1:200 diluted duplex oligo, 1XTango buffer (Thermo Fisher Scientific#BY5), 50mM DTT, 5mM ATP, 1U BbsI (New England Biolabs#R3539) and 0.5U T4 DNA ligase. The reaction was incubated in 6 cycles of 37°C for 5’ followed by 21°C for 5’. Plasmids were transformed into competent bacteria and prepped as described above. Sequences were checked by Sanger sequencing.

#### Transfection

5x10^5^ mESCs in 1ml of media were seeded in 0.2% gelatin coated wells of a 6-well plate and incubated for 6h at 37°C. For transfection 1.25µg of each plasmid were incubated together with 250µl OptiMEM and 7.5µl *Trans*IT-X2® for 20’’ at room temperature. After that, the reaction was added dropwise to the mESCs. After 24h at 37°C MEFs were added to the stem cells again. To select positive clones, 48h post transfection ESC media was supplemented with 1µg/ml Puromycin (Thermo Fisher Scientific#A1113803). After 4 days, selection media was replaced by normal ESC media again for recovery.

#### Genotyping-Cell lines

Single colonies were picked into a 96-well plate and analyzed for deletions by PCR analysis and Sanger sequencing. For genotyping, 15µl of cells were lysed in 35µl lysis buffer containing 3.5µg Proteinase K (Thermo Fisher Scientific#BP1700-100). The reaction was incubated for 2’ at 37°C, 2h at 55°C and finally for 10’ at 95°C. To identify clones that carried the deletion, lysed cells were taken as template for PCR. Each 20µl PCR reaction contained 1XREDTaq® ReadyMix™ PCR Reaction Mix (Sigma-Aldrich#R2648), 10µM of each primer and 2µl of the lysis mix as a template. The PCR program was 95°C for 2’, followed by 35 cycles of 95°C for 30’’, 60°C for 30’’, 72°C for 40’’ and a 2’ extension cycle at 72°C. The PCR products were visualized by agarose gel electrophoresis in TBE buffer and stained with SafeView.

sgRNAs, ssODN, homology arms and genotyping primer sequences for generating the IG-TRE deletion and the SOX2-binding site deletion within the IG-TRE are listed in Table S2.

### RNA and DNA extraction

DNA/RNA extraction was performed with the AllPrep DNA/RNA Mini and Micro Kits (Qiagen#80204, 80284) according to the manufactor’s instructions.

### Pyro-Sequencing Methylation analysis

The procedure and primers for this DNA methylation analysis were described previously^44–46^. In brief, 1μg DNA was treated using the EZ-96 DNA methylation Kit (Zymo Research#D5032) in accordance with the manufacturer’s instruction. Bisulfite-treated DNA was eluted in 30μl of elution buffer. Amplicons were generated in a 25μl reaction volume containing 100nM forward and reverse primers, 1.25U of HotstarTaq DNA Polymerase (Qiagen#203203), 0.2mM dNTPs, and 5μl of bisulfite-treated DNA. PCR cycle conditions consisted of an initial activation step of 95°C for 15’, followed by 50 cycles of 94°C for 30’’, specific annealing temperature for 30’’, and extension at 72°C for 30’’, followed by a final extension at 72°C for 10’ (primers are listed in Table S3). Pyrosequencing was carried on a PSQ HS96 System using PyroMark Gold Q96 SQA Reagents (Qiagen#972812). The degree of methylation at CpG sites (without distinguishing between maternal and paternal alleles) was determined by the pyro-Q CpG software.

### Reverse transcription and quantitative real-time PCR (qPCR)

RNA was treated with RNase-free DNaseI (Thermo Fisher Scientific#EN0521) prior to cDNA synthesis using RevertAid H Minus First Strand cDNA Synthesis Kit with oligo dT and random hexamer primers (Thermo Fisher Scientific#K1632). qPCR was performed with BrilliantII SYBR® Green QPCR Master Mix (Agilent Technologies#600882) in a 384-well plate on a QuantStudio™ 5 Real-Time PCR System (Applied Biosystems#A34322). Relative quantification of gene expression was normalized to the geometrical mean of GAPDH and β-Actin expression levels (primers are listed in Table S4) and calculated using the ΔΔCT method, plotted as 2^-ΔΔCT^. Pyrosequencing was carried on a PSQ HS96 System using PyroMark Gold Q96 SQA Reagents (Qiagen#972812). SNP percentage was determined by the pyromark^TM^ MD 1.0 software.

### Chromatin immunoprecipitation (ChIP)-qPCR

ChIP was performed as previously described with some modifications^47,48^. 5x10^6^ mESCs or 100mg brain tissue were crosslinked in 1% formaldehyde for 10’ and subsequently quenched with Tris pH 8.0 (250mM final) for 10’. The quenched cells were washed twice with ice-cold PBS supplemented with a protease inhibitor cocktail (EDTA-free cOmplete™, Sigma-Aldrich#11873580001), flash frozen in liquid nitrogen, and stored at -80°C. Crosslinked cells were thawed on ice and then lysed sequentially on ice and for 10’ at each step in each of the following buffers: LB1 (50mM HEPES-KOH pH 7.4, 140mM NaCl, 1mM EDTA, 0.5mM EGTA, 10% Glycerol, 0.5% NP-40, 0.25% Triton-X-100 and EDTA-free cOmplete™), LB2 (10mM Tris-HCl pH 8.0, 200mM NaCl, 1mM EDTA, 0.5mM EGTA and EDTA-free cOmplete™), and SDS-shearing lysis buffer (10mM Tris-HCl pH 8, 1mM EDTA, 0.15% SDS and EDTA-free cOmplete™). Nuclei from brain lysates were counted with Trypan Blue staining and 5x10^6^ cells were taken forward for each ChIP. The lysates were sonicated (Bioruptor Plus, Diagenode) at 4°C to generate DNA fragments of 100-500bp (2 repetitions of 10 sonication cycles of 30 seconds on and 30 seconds off) and the sonicated lysates were subsequently clarified by centrifugation (15,000rpm for 10’ at 4°C). 10% of the mixture was taken as input control. The sonicated lysate was topped up to 1mL with LB3-500NaCl (20mM Tris-HCl pH 7.5, 500mM NaCl, 1mM EDTA, 0.5mM EGTA, 1% Triton-X-100, 0.1% Sodium deoxycholate, 0.1% SDS and EDTA-free cOmplete™). This mixture was incubated overnight at 4°C with Protein G Dynabeads (Invitrogen#10004D) that had been pre-blocked with 0.5% BSA and mixed with 5µg Goat polyclonal SOX2 antibody (R&D Systems#AF2018) or Protein A Dynabeads (Invitrogen#10002D) mixed with 5µg Rabbit polyclonal Zfp281/ZNF281 antibody (abcam#ab101318). To remove non-specifically bound proteins from the ChIP beads were washed five times with RIPA Buffer (50mM HEPES-KOH pH 7.4, 500mM LiCl, 1mM EDTA, 1% NP-40 and 0.7% Sodium deoxycholate) and once with TE Buffer (50mM Tris-HCl pH 8.0, 10mM EDTA) at 4°C. The DNA-protein complex was eluted from the beads in Elution Buffer (50mM Tris-HCl, pH 8.0, 10mM EDTA and 1% SDS) for 20’ at 65°C, and reverse-crosslinked overnight at 65°C. The eluted samples were then treated with 10mg/ml RNase A (Sigma-Aldrich#R4875) and 20mg/ml Proteinase K, and purified using the QIAquick PCR Purification Kit.

qPCR was performed with BrilliantII SYBR® Green qPCR Master Mix (Agilent Technologies#600882) in a 384-well plate on a QuantStudio™5 Real-Time PCR System (Applied Biosystems#A34322). The ChIP-qPCR fold enrichments were calculated according to the percent input method, where signals obtained from the ChIP are divided by signals obtained from the input sample. qPCR primers are listed in Table S5.

### CRISPR-zygote injections and generation of IG-TRE^Δ550^ knockout mice

Deletion of 550bp encompassing the distal region of a germline-derived intergenic differentially methylated region (IG-DMR, MGI:3051678) including the transcriptional regulation element (IG-TRE) was made by CRISPR/Cas9 gene editing by the Mary Lyon Centre at MRC Harwell as part of the Genome Editing Mice for Medicine (GEMM) program. Constructs were delivered by electroporation into 1-cell stage embryos from a C57BL/6J genetic background. Briefly, 90:10 deactivated Cas9:Cas9 protein, sgRNAs and ssODNs were diluted and mixed in Electroporation buffer (EB; *Gibco* Opti-MEM I Reduced Serum Media (Thermo Fisher Scientific) to the working concentrations of 650ng/μl, 130ng/μl total and 400ng/μl, respectively. Embryos were electroporated using the following conditions: 30V, 3ms pulse length, 100ms pulse interval, 12 pulses. Electroporated embryos were re-implanted in CD1 pseudo-pregnant females. Host females were allowed to litter and rear F0 progeny.

For genotyping, genomic DNA was extracted from ear clip biopsies and amplified in a PCR reaction using the ThermoFisher SuperFi II PCR kit under the following conditions: 60^ᵒ^C Annealing Temperature, 1.25’ Elongation time, wildtype product size: 1412bp, Mutant product size: 862bp. All amplicons were sent for Sanger sequencing to check for integration of the donor oligo sequence at the target site. No off-target activity was detected in the animals.

Copy counting of the donor sequence was carried out by ddPCR at the F1 stage to confirm donor oligos were inserted once on target into the genome. A Taqman assay was used to copy count the donor sequence compared against a VIC-labelled reference assay for Dot1l. This ddPCR assay is specific to the wildtype allele and only wildtype alleles are expected to be recognised by this assay. Therefore, wildtype controls are expected to call at 2 copies and a single integration for a correct mutation is expected to call at 1 copy for F1 (heterozygote) animals. No additional donor integrations were detected in the animals taken forward to establish the colony.

sgRNAs, ssODN, off target primers, taqman assay primers, taqman assay probes and genotyping primer sequences for generating the IG-TRE^Δ550^ deletion in mice are listed in Table S6.

### Mice lines

CAST/EiJ (RRID:IMSR_JAX:000928) and IG-TRE^Δ550^ knockout mice were obtained from MRC Harwell Institute, and maintained under a 12h light-dark cycle at 22°C degrees (+/-2°C) and 55% humidity (+/-10%). Mice were monitored for health and activity and were given ad libitum access to water and standard mouse chow. For pure and hybrid breeding experiments, mice were mated at 8-12 weeks of age. Male and female heterozygote F1 mice carrying the IG-TRE^Δ550^ deletion were crossed with wildtype CAST/EiJ, wildtype C57BL/6J or IG-CGI deletion partners, and F2 animals harboring the deletion allele were analyzed at different stages. All animal experiments were performed in accordance with the Animals (Scientific Procedures) Act 1986 Amendment Regulations 2012 following ethical review by the University of Cambridge Animal Welfare and Ethical Review Body. Animal experiments were approved by the UK Home Office project license #PP8193772. All efforts were made to minimize animal discomfort.

### Genotyping-Mice

For genotyping, DNA was extracted from ear clips using a solution containing NaOH 1M and EDTA 0.5M pH8.0 in DDW, incubated for 1h at 95°C and neutralized in Tris-HCl 40Mm pH5.0. Alternatively, DNA was isolated by lysing cells in lysis buffer (0.1M Tris buffer, 0.2M NaCl, 0.005M EDTA, 0.2% SDS) with 10mg/ml Proteinease K at 55°C, precipitated with Iso-Propanol, washed with 70% Ethanol and resuspended in TE buffer. Genotyping of mouse strains and alleles was done by PCR, primers are listed in Tables S2 and S6.

### Tissue and Embryo analysis

Animals harboring the IG-TRE^Δ550^ deletion allele were analyzed at different ages. At 9 weeks, measurements of adult animal weight (g), brain weight (mg), liver weight (g), fat weight (mg) and animal length (cm) were recorded. Brains were collected for expression analysis. At embryonic day E15.5 measurements of embryo and placenta lengths (cm) were recorded and brain, liver and placenta were collected for expression analysis. At E8.5 the whole embryo was used for bulk DNA and RNA purification and analysis.

### Image analysis

Measurements of embryo and placenta lengths at E15.5 were carried out with ImageJ software and calculated as Feret’s Diameter.

### Statistical analysis

At least three biological replicates were performed for all experiments. Statistical differences were determined using a two-tailed unpaired Student’s t-test. Data are shown as means with error bars representing the standard error. P-values of less than 0.05 were considered significant.

## Data availability

All relevant data and details of resources can be found within the article and its supplementary information.

## Acknowledgements

This work was supported by an MRC programme grant (MR/X018407/1) awarded to A.C.F-S. A.W-S is supported by the Human Frontier Science Program (LT000966/2020-L), the Rothschild Yad Hanadiv Fellowship program, the Humanitarian trust, the 2020 Israel National Postdoctoral Award for Advancing Women in Science and the 2024 Weizmann Institute of Science Women’s Postdoctoral Career Development Award in Science. We thank Anjuli Freeman for mouse colony husbandry and the Ferguson-Smith group members for their discussion and advice. The IG-TRE^Δ550^ mice used in this study were obtained from the Mary Lyon Centre at MRC Harwell and the following award is acknowledged: MC_UP_2201/2. Figures were created with BioRender.com.

## Author contributions

A.W-S and A.C.F-S. conceived and designed the experiments, performed data analysis, and its interpretation. A.W-S, F.L.D, A.M-B, L.T.N.N, M.M and R.C.R carried out experiments. C.A.E assisted with interpreting the results and gave input on the manuscript text. A.W-S prepared the figures. A.W-S and A.F-S wrote the manuscript.

## Materials & Correspondence

All relevant data and details of resources can be found within the article and its supplementary information. Correspondence and material requests should be addressed to.

