## Supplemental information for "Regulatory Mechanisms of Maternal Imprinting at the Dlk1-Dio3 Domain"

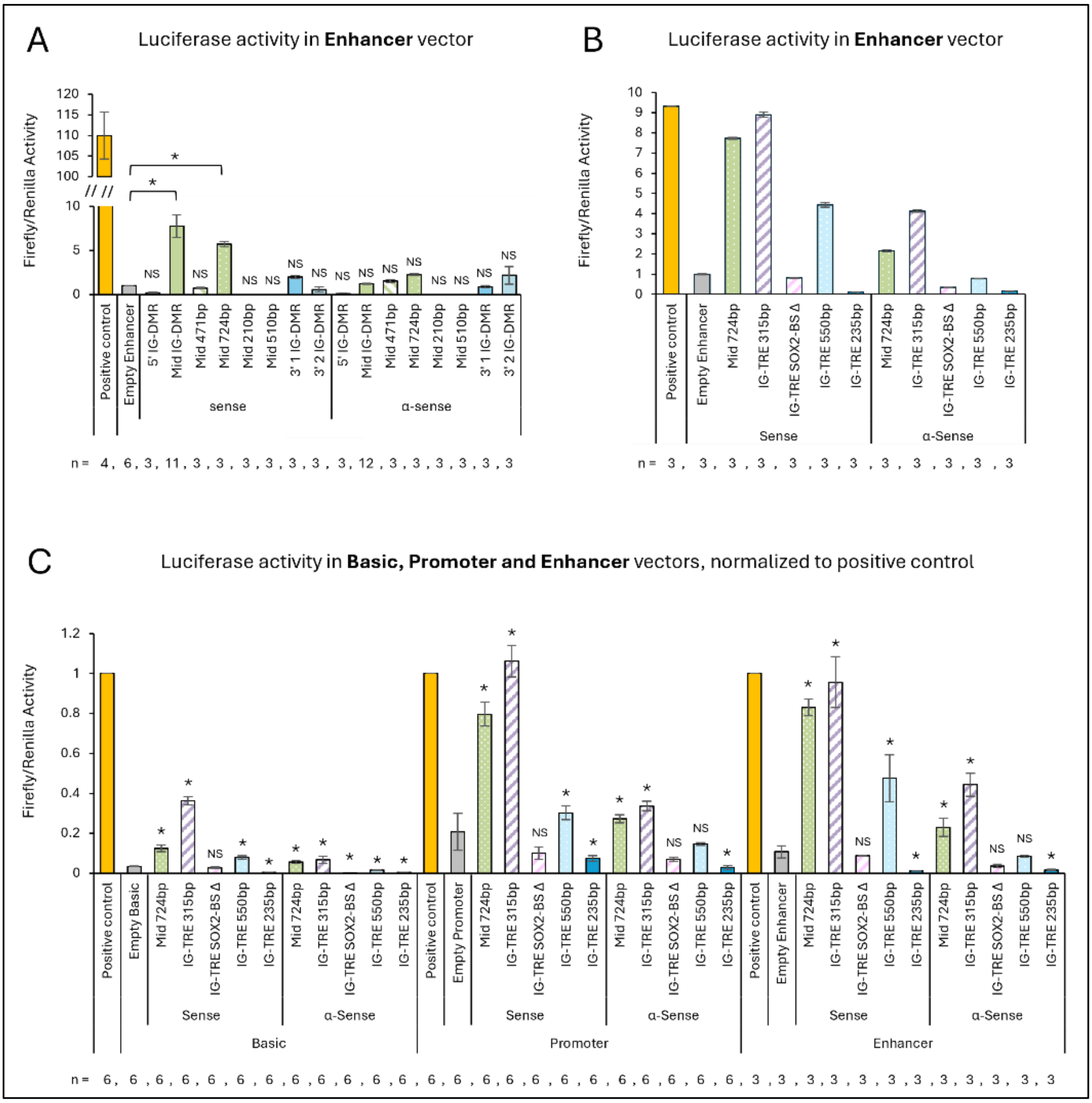


**Figure S1- Enhancer and silencer activity detected at the IG-TRE. A+B.** Graphical representation of the luciferase activity in the pGL3 Enhancer vector, normalized to the empty enhancer vector. **C**. Graphical representation of the luciferase activity in the pGL3 Enhancer vector normalized to the positive control vector. The Mid 724bp fragment shows activity in sense orientation only, suggesting promoter activity. The Mid 315bp fragment shows activity similar to the 724bp fragment, suggesting enhancer activity. This activity is lost upon deletion of the SOX2-binding site from within the 315bp fragment. In parallel, the 235bp fragment, downstream of the 315bp region, shows silencing activity, indicating dual activity in a minimal region of 550bp in the IG-TRE with both enhancing and silencing regulation. Each construct was measured twice in triplicates wells. The firefly luciferase values were normalized to the *Renilla* values and then each test construct was normalized to the corresponding empty vector. NS- not significant. Asterisks indicate statistical significance in comparison to the empty vector using a two-tailed unpaired Student's t-test.

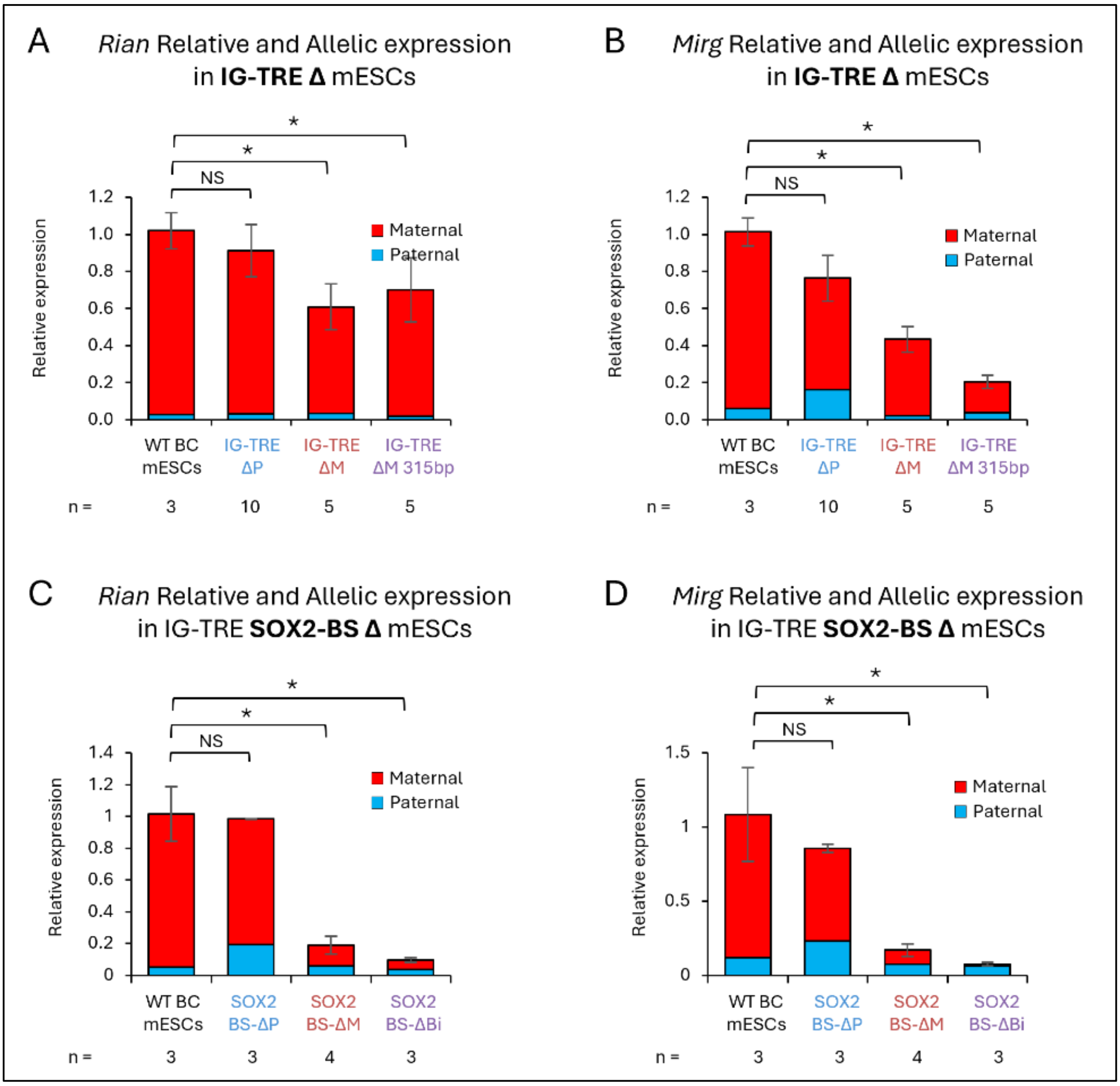


**Figure S2- Deleting the entire maternal IG-TRE or even just the SOX2-binding site *in vitro* results in dysregulation of the maternal transcripts *Rian* and *Mirg*. A.** Graphical representation of *Rian* expression in mESC clones deleted for the entire IG-TRE (1.3kb) shows loss of *Rian* expression upon maternal or biallelic deletions. **B.** Graphical representation of *Mirg* expression in mESC clones deleted for the entire IG-TRE (1.3kb) shows loss of *Mirg* expression upon maternal or biallelic deletions. **C.** Graphical representation of *Rian* expression in mESC clones deleted for the SOX2-binding site within the 315bp of the IG-TRE shows loss of *Rian* expression upon maternal or biallelic deletions. **D.** Graphical representation of *Mirg* expression in mESC clones deleted for the SOX2-binding site within the 315bp of the IG-TRE shows loss of *Mirg* expression upon maternal or biallelic deletions. NS- not significant. Asterisks indicate statistical significance in comparison to wildtype using a two-tailed unpaired Student's t-test.

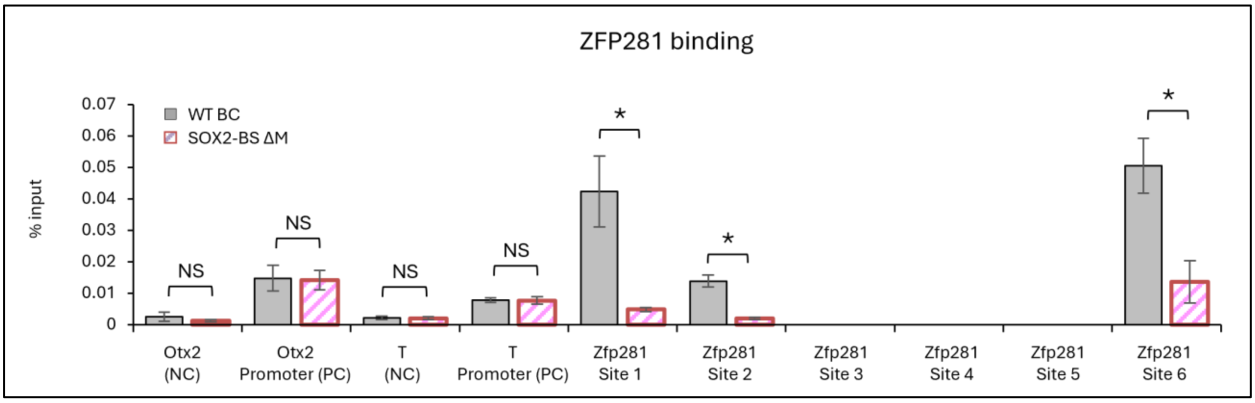


**Figure S3- ZFP281 binding is dependent on Sox2 binding on the maternal chromosome.** Graphical representation of ZFP281 binding at the IG-TRE, determined by ChIP-qPCR, shows depletion of ZFP281 binding at binding sites 1, 2 and 6, when the SOX2-binding site is deleted from the 315bp region. No binding was observed at predicted binding sites 3, 4 and 5 and no change in binding was observed at control sites and the *Otx2* and *T* genes. NS- not significant. Asterisks indicate statistical significance in comparison to wildtype using a two-tailed unpaired Student's t-test.

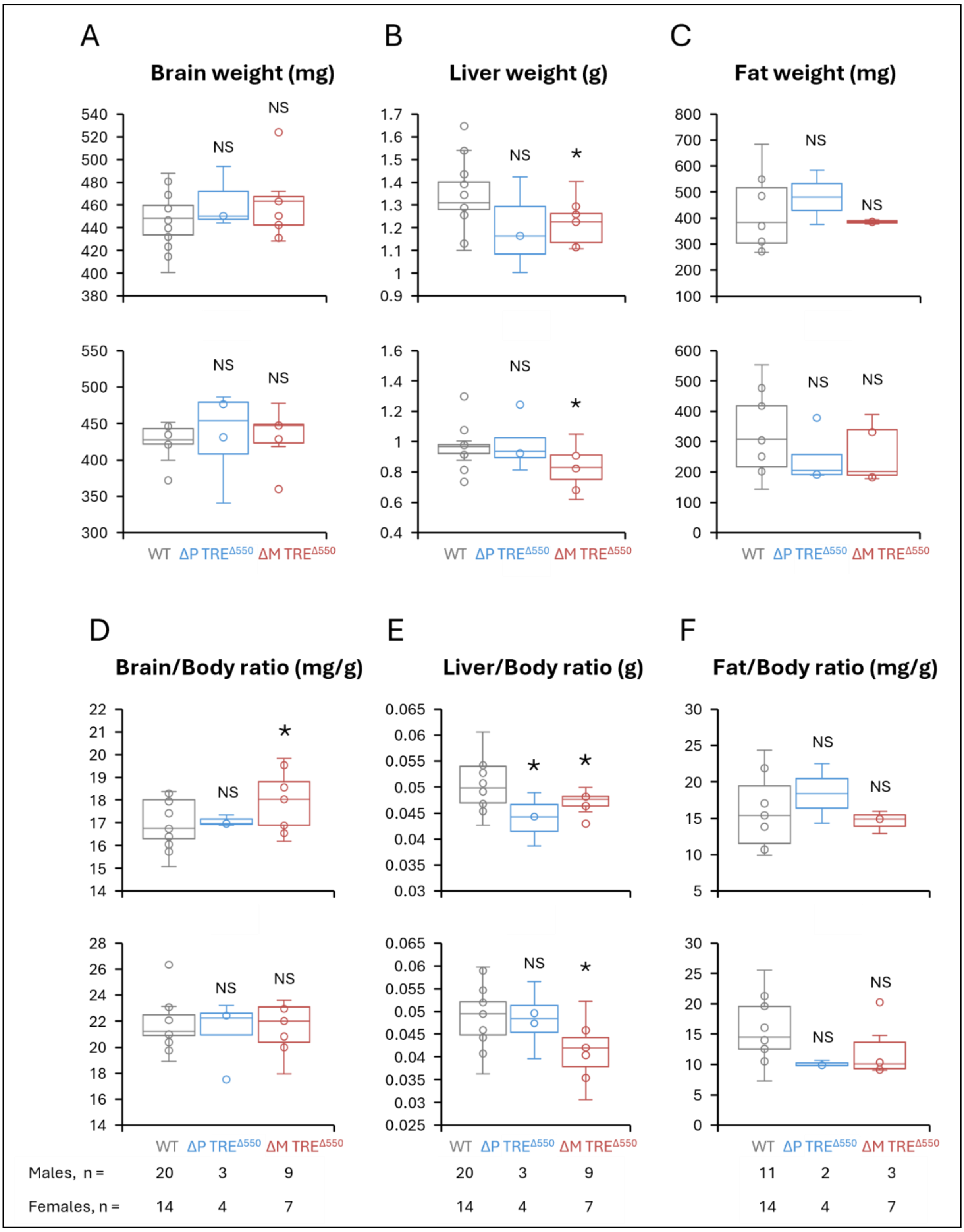


**Figure S4- Deletion of the IG-TRE$^{\Delta550}$ *in vivo* is viable with a slight reduction in adult liver size. A-F.** Box plot representation of brain weight (mg), liver weight (g) and fat weight (mg) of 9 week-old adult animals. $N_{WT}$=20, $N_{\Delta P\text{-}TRE}{}^{\Delta550}$=3, $N_{\Delta M\text{-}TRE}{}^{\Delta550}$=9 biologically independent males were measured for brain weight, liver weight, brain/body ratio and liver/body ratio; and $N_{WT}$=11, $N_{\Delta P\text{-}TRE}{}^{\Delta550}$=2, $N_{\Delta M\text{-}TRE}{}^{\Delta550}$=3 biologically independent males were measured for fat weight and fat/body ratio. $N_{WT}$=14, $N_{\Delta P\text{-}TRE}{}^{\Delta550}$=4, $N_{\Delta M\text{-}TRE}{}^{\Delta550}$=7 biologically independent females were measured for all categories. NS- not significant. Asterisks indicate statistical significance in comparison to wildtype using a two-tailed unpaired Student's t-test.

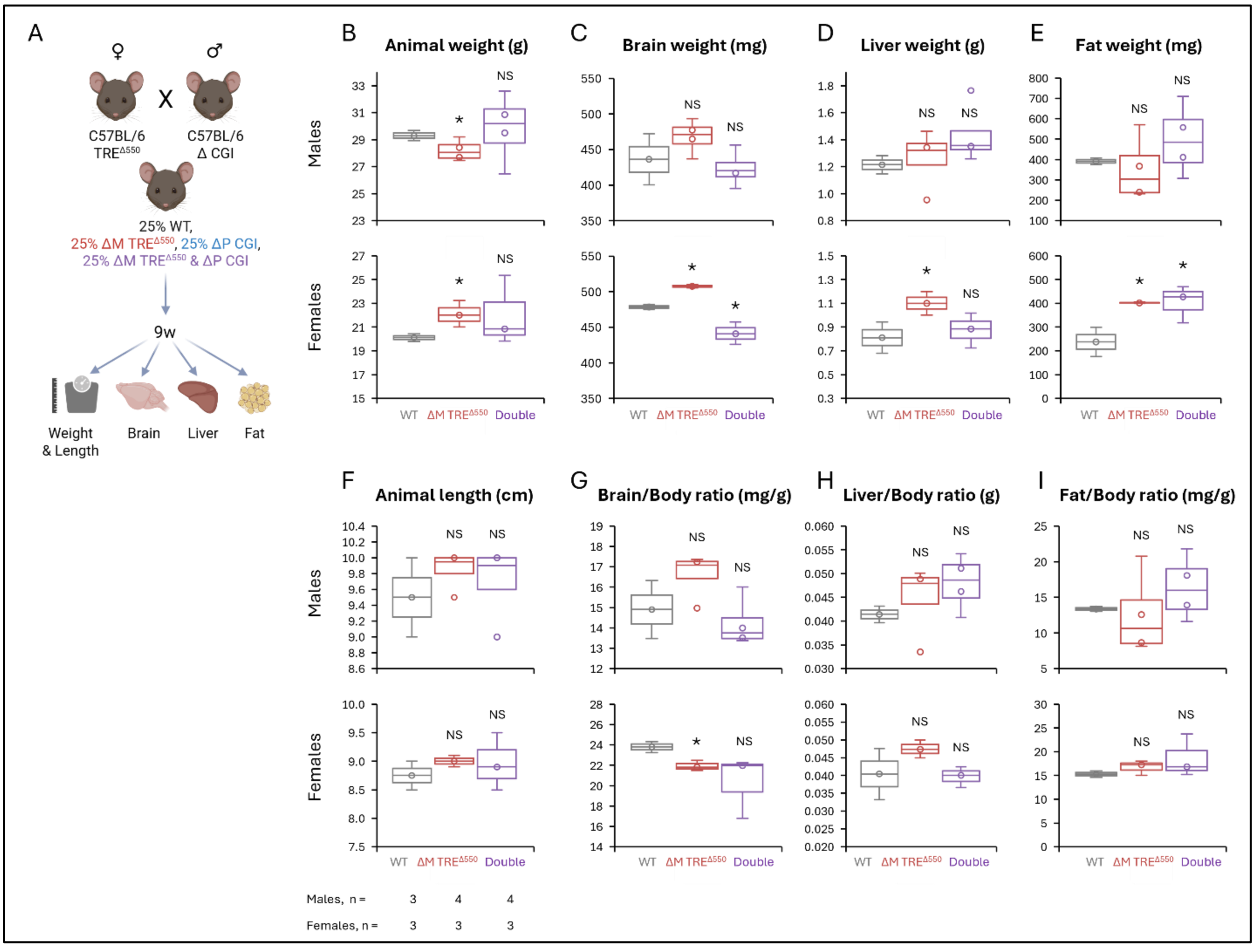


**Figure S5- IG-CGI$^{\Delta P}$ and IG-TRE$^{\Delta 550}$ double deletion embryos are normal in size and rescue the lethality of the paternal IG-CGI deletion *in vivo*. A.** Schematic representation of the predicted outcome of crossing IG-CGI$^{\Delta P}$ and IG-TRE$^{\Delta 550}$ mouse strains. In each litter, 25% of the pups are wildtype littermates (pure C57BL/6) and kept as controls. Tissues were collected for expression at 9 weeks. **B-I.** Box plot representation of total body weight (g), brain weight (mg), liver weight (g), fat weight (mg) and body length (cm) of 9 week-old adult animals. $N_{WT}$=3, $N_{\Delta M\text{-}TRE}{}^{\Delta 550}$=4 and $N_{\Delta M\text{-}TRE}{}^{\Delta 550}{}_{\&\Delta P\text{-}CGI}$=4 biologically independent males and $N_{WT}$=3, $N_{\Delta M\text{-}TRE}{}^{\Delta 550}$=3 and $N_{\Delta M\text{-}TRE}{}^{\Delta 550}{}_{\&\Delta P\text{-}CGI}$=3 biologically independent females. NS- not significant. Asterisks indicate statistical significance in comparison to wildtype using a two-tailed unpaired Student's t-test.

**Table S1**- primers used to generate and sequence luciferase constructs

| Fragment | Primer | Sequence 5’ → 3’ |
|---|---|---|
| IG-DMR 5' | SF8 (SacI) forward | CACGGCACAGAGCAGTAGCG |
| | SR9 (SacI) reverse | ACTGCCATTTAGAATTTGAG |
| IG-DMR Mid | IG-DMR Mid 1F NheI forward | AACTACCGCTACGGTTCATA |
| | IG-DMR Mid 1R NheI reverse | CATCCCACGAGGAACTGGT |
| | Mid forward | AACTACCGCTACGGTTCATA |
| | Mid NheI reverse | GGCCGCTAGCCATCCCACGAGGAACTGGT |
| | Mid KpnI reverse | GGCCGGTACCCATCCCACGAGGAACTGGT |
| IG-DMR 3'1 | IGDMR 3’1F NheI forward | CCTTAGGGCTTCCAAGC |
| | IGDMR 3’1R NheI reverse | CTGGCGAGCAACAGTGAA |
| IG-DMR 3'2 | IGDMR 3’2F NheI forward | GCTTCACTGTTGCTCGCC |
| | IGDMR 3’2R NheI reverse | CATGCTCAGAGCCAGCAG |
| 315bp | NheI forward (also used for 550bp) | GGCCGCTAGCCGGTCTCGGAGTTCTTGTC |
| | NheI reverse | GGCCGCTAGCAGACGAGTTGGCAGGACG |
| | KpnI forward (also used for 550bp) | GGCCGGTACCCGGTCTCGGAGTTCTTGTC |
| | KpnI reverse | GGCCGGTACCAGACGAGTTGGCAGGACG |
| 550bp | NheI reverse (also used for 235bp) | GGCCGCTAGCTGGTCAACAAAAGCATCTCTGT |
| | KpnI reverse (also used for 235bp) | GGCCGGTACCTGGTCAACAAAAGCATCTCTGT |
| 235bp | NheI forward | GGCCGCTAGCGAGCCCTGTGGTTAACTAAGC |
| | KpnI forward | GGCCGGTACCGAGCCCTGTGGTTAACTAAGC |
| pGL3 empty vector | RV primer forward | CTAGCAAAATAGGCTGTCCC |
| | GL primer reverse | TATGTTTTTGGCGTCTTCCA |
| pGL3 promoter vector | Forward | AGGTACCGAGCTCTTACGC |
| | Reverse | GCCGGGCCTTTCTTTATGTT |
| pGL3 enhancer vector | Forward | GGGAGGTGTGGGAGGTTTT |
| | Reverse | AAGGAGCTGACTGGGTTGAA |

**Table S2**- gRNAs and genotyping primers for CRISPR in mESCs

| Target | Primer | Sequence 5’ → 3’ |
|---|---|---|
| 5’ IG-TRE | gRNA 1 forward | CACCGCTGTAGCCTTGCTAGACTG |
| | gRNA 1 reverse | AAACCAGTCTAGCAAGGCTACAGC |
| 3’ IG-TRE | gRNA 2 forward | CACCgCCATGGCTTACAAGCTACTG |
| | gRNA 2 reverse | AAACCAGTAGCTTGTAAGCCATGGC |
| SOX2-BS | gRNA forward | CACCgAGAAGGGATGAGACTCCTCT |
| | gRNA reverse | AAACAGAGGAGTCTCATCCCTTCTC |
| | single-stranded oligo donor (ssODN) | ACGGCTCTTTCCCAGAATCCACTGGGCCTGTTTTGGGGGCAGCTTA GTTAACCACAGGGCTCTGGTTAGAAGGGATGAtACgCCTCTAGGATT CCCAGAGTTTCTCGGTCTTCT |
| IG-TRE Δ | Genotyping forward | TGACTTCCTTCAGCCACAGT |
| | Genotyping reverse | GGCAAACCCCAACTTAGCAA |
| | Genotyping internal reverse | CAGGATGCCAAGGGTTGTG |
| SOX2-BS Δ | Genotyping forward | CGGTCTCGGAGTTCTTGTC |
| | Genotyping reverse | AGACGAGTTGGCAGGACG |

**Table S3**- primers for bisulfite sequencing

| Target | Primer | Sequence 5’ → 3’ |
|---|---|---|
| IG-CGI<br>With biotin for qPCR and pyro methylation analysis | Forward | GTGGTTTGTTATGGGTAAGTTT |
| | Reverse | CCCTTCCCTCACTCCAAAAATTAA |
| | Sequencing | TGGTTTATTGTATATAATGT |
| IG-TRE<br>With biotin for qPCR and pyro methylation analysis | Forward | GTTGGGGTTTGTTAGTTATTTATATGTTAT |
| | Reverse | AAAACATACTCTCCACTATAACTAATT |
| | Sequencing | CTATAACTAATTACAACACCAC |
| Gtl2-DMR<br>With biotin for qPCR and pyro methylation analysis | Forward | AGTTATTTTTTGTTTGAAAGGATGTGTA |
| | Reverse | CTAACTTTAAAAAAAAATCCCCAACACT |
| | Sequencing | GAAAGGATGTGTAAAAATGA |

**Table S4**- primers for qPCR for gene expression

| Target | Primer | Sequence 5’ → 3’ |
|---|---|---|
| *GAPDH*- housekeeping gene (ex2F+3R) | Forward | AGGTCGGTGTGAACGGATTTG |
| | Reverse | TGTAGACCATGTAGTTGAGGTCA |
| *B2M*- housekeeping gene (ex1F+2R) | Forward | GGCTGTATTCCCCTCCATCG |
| | Reverse | CCAGTTGGTAACAATGCCATGT |
| *Dlk1*- ex5<br>With biotin for qPCR and pyro SNP analysis | Forward | CGCAAGAAGAAGAACCTCCTGT |
| | Reverse | ACGCTGCTTAGATCTCCTCATCA |
| | Sequence | CAGCCTCCTTGTTGAA |
| *Gtl2*- ex10<br>With biotin for qPCR and pyro SNP analysis | Forward | AGCCACCTATTTACAAATGGACTC |
| | Reverse | CATCCCCATGAGAAACCTGTT |
| | Sequence | CATAGAGACACAAACATAGT |
| *Mirg*- ex16<br>With biotin for qPCR and pyro SNP analysis | Forward | CTCAGGAGCGGATGTTCAAG |
| | Reverse | GATGTCCCCAGTGGAATGTC |
| | Sequence | GGAACCCTGCCTATG |
| *Rian*- ex4<br>With biotin for qPCR and pyro SNP analysis | Forward | GAGACCTTGGCAGTGACCG |
| | Reverse | CCTGGCCCAAAAAGCCTC |
| | Sequence | CTTGGCAGTGACCGC |
| *Dio3*- ex4<br>With biotin for qPCR and pyro SNP analysis | Forward | GAGGGATGCGAGAACTTTTTG |
| | Reverse | GCGCTTTCTGCCTAGGACT |
| | Sequence | TTTTGGAGAAGGGATT |

**Table S5**- primers for ChIP-qPCR

| Target | Primer | Sequence 5’ → 3’ |
|---|---|---|
| SOX2 Binding site at IG-TRE | Forward | TCCAATCTTCAAACACCTCCTG |
| | Reverse | TCCACTGGGCCTGTTTTGG |
| SOX2 Binding site at IG-TRE<br>With biotin for qPCR and pyro SNP analysis | Forward | CCAATCTTCAAACACCTCCTG |
| | Reverse | CCACTGGGCCTGTTTTGG |
| | Sequence | CAAAACAATAAGAAGGGA |
| SOX2 Binding at Scmh1- positive control<br>(from: Boumahdi *et al*. PMID: 24909994) | Forward | AGCCAACAACGGCACTAAGA |
| | Reverse | CATGGAAGGATGGAGTGGGT |
| SOX2 Binding at Ceacam1- negative control<br>(from: Boumahdi *et al*. PMID: 24909994) | Forward | CTGAAGAGTGGATGGTAAGG |
| | Reverse | CACACCGCAAGGTCAGAATG |
| ZFP281 Binding site #1a at TRE<br>With biotin for qPCR and pyro SNP analysis | Forward | CCAATAGCATGGAGCTCAGGTTAG |
| | Reverse | CGAGACCGGTGTTGTAAGTGC |
| | Sequence | GTGCATGCAGGATCGC |

| | | |
|---|---|---|
| ZFP281 Binding site #1b at TRE<br>With biotin for qPCR and pyro SNP analysis | Forward | AGTGAGGGAAGGGCTGCATTA |
| | Reverse | GCTAACCTGAGCTCCATGCTATTG |
| | Sequence | ATGCCTTGAGCACAG |
| ZFP281 Binding site #2 at TRE<br>With biotin for qPCR and pyro SNP analysis | Forward | TGACTTCCTTCAGCCACAGTTC |
| | Reverse | GATGCTCAGAAAGGCAGTGG |
| | Sequence | GTGCATGCAGGATCGC |
| ZFP281 Binding site #3 downstream of TRE<br>With biotin for qPCR and pyro SNP analysis | Forward | TGATTCACCACAATTCGTTCTG |
| | Reverse | CGGTCATCAATTTGCCATTTCT |
| | Sequence | CGTTCTGGTTTGTGATTA |
| ZFP281 Binding site #4 upstream of Gtl2<br>With biotin for qPCR and pyro SNP analysis | Forward | TTCGTGTTAGCGAGACAACATT |
| | Reverse | CCACCCCCAGGAGAAAGTATATC |
| | Sequence | TGTTAGCGAGACAACATT |
| ZFP281 Binding site #5 at Gtl2-DMR<br>With biotin for qPCR and pyro SNP analysis | Forward | ATGGGGGTGCATAGCGAC |
| | Reverse | AGACTCCAATAGCCCAACCACC |
| | Sequence | GCCCAACCACCTGAG |
| ZFP281 Binding site #6 at Gtl2-DMR | Forward | TCATCTGTACCCCTCCCATTG |
| | Reverse | AGGGAAATGGGTAGGGGC |
| ZFP281 Binding at Otx2 promoter- positive control<br>(from: Huang *et al*. PMID: 29168693) | Forward | CCCGAGGAAAGCAATTACAA |
| | Reverse | AAATGGCCCCAATCAAGTTT |
| ZFP281 Binding at Otx2- negative control<br>(from: Huang *et al*. PMID: 29168693) | Forward | AATGCCTGGCTAAAACTGGA |
| | Reverse | CTTCAATGCTGACTGCTTGG |
| ZFP281 Binding at T promoter- positive control<br>(from: Huang *et al*. PMID: 29168693) | Forward | GCTTCAAGGAGCTAACTAACGAG |
| | Reverse | CCAGCAAGAAAGAGTACATGGC |
| ZFP281 Binding at T- negative control<br>(from: Huang *et al*. PMID: 29168693) | Forward | CCTAGCTGTTTTGGCTTTGG |
| | Reverse | GCTGAGGCAGGAGAATCAAG |

### **Table S6**- gRNAs and genotyping primers for CRISPR in mice

| Target | Primer | Sequence 5' → 3' |
|---|---|---|
| 5' IG-TRE | sgRNA 1 | CAGTCTAGCAAGGCTACAGC, PAM: AGG |
| | sgRNA 2 | ATATAGTCCACAGTCTAGCA, PAM: AGG |
| 3' IG-TRE | sgRNA 3 | ATATAGTCCACAGTCTAGCA, PAM: TGG |
| | sgRNA 4 | TTGGTTTACCAGTTCCTCGT, PAM: GGG |
| single-stranded oligo donor (ssODN) | CCTTCAGCCACAGTTCCCAGGCAGCCCTTGGCTGGAGGCATTGCGATCCTGCATGCACTTACAACACCGGTCTCGGAGTTCTTGTCTCCCTGCTGTAGCCGTGTCTGCTGCAGTGTGCCAGTTTTTTCGTGGTACACAGAGATGCTTTTGTTGACCACAACCCTTGGCATCCTGGCTCACCGTTGTCTAGAAACACTTTT | |
| IG-TRE Δ550 | Genotyping forward | GCCGCTATGCTATGCTGTTTC |
| | Genotyping reverse | GTGCCTCAGTGCCGTGTTAG |
| | Sequencing forward | CCCAGGACCTCCAATAGCAT |
| Off target 1:<br>chr12:54356165-54356187 (Intergenic)<br>ATA**C**AG**G**C**T**ACAGTCTAGCA TGG | Off target 1 forward | TACAGAGGGACTCTGGTATCTACA |
| | Off target 1 reverse | GGGATTTGTGGGTCTCGCA |
| | Off target 1 sequence forward | TCTACATGCAAAAGATGAACTGC |
| Off target 2:<br>chr12:109960424-109960446 (Intergenic)<br>A**A**A**G**AGT**T**C**C**CAGTCTAGCA GGG | Off target 2 forward | ATGACCTCAGTCTGCAAGGC |
| | Off target 2 reverse | TCCAGGGCAGGCATTACAAG |
| | Off target 2 sequence forward | ACCTGGAGTGGTGAGCCTAT |
| IG-TRE copy count taqman assay | Forward | CCCAGGTTGCCCTTAGTAAAT |
| | Reverse | CCAGAGTTTCTCGGTCTTCTTT |
| | Probe (FAM) | TGCGGGAGACACCCTGTTTCTTTA |
| *Dot1l* copy count reference taqman assay | Forward | GCCCCAGCACGACCATT |
| | Reverse | TAGTTGGCATCCTTATGCTTCATC |
| | Probe (VIC) | CCCAACAGGCCTGGATTCTCAATGC |
